# A discrete Connexin26+ neural crest lineage emerges in mid-life and mediates enhanced central brainstem responses to elevated CO_2_ levels for deep breathing

**DOI:** 10.64898/2026.09.16.752084

**Authors:** Xintao Zhang, Jie Zhang, John M. J. Lapage, Alexander V. Gourine, Nicholas Dale, Georgy Koentges

**Affiliations:** University of Warwick, School of Life Sciences, Coventry CV4 7AL, UK; Warwick Neuroscience Group, University College London, London, UK; Laboratory of Systems Biomedicine and Evolution, University College London, London, UK; Centre for Cardiovascular and Metabolic Neuroscience, Department of Neuroscience, Physiology and Pharmacology, University College London, London, UK

## Abstract

The precise control of breathing is fundamental to vertebrate survival. Increased PCO_2_ in blood and brain parenchyma causes an increase in both the frequency and volume of lung ventilation. We have previously demonstrated that CO_2_ directly binds to connexin26 (Cx26) hemichannels causing them to open and allow release ATP. We now document the role of Cx26 as a direct physiological CO_2_ sensor *in vivo*. Here we describe a unique Cx26+ neural crest cell lineage that populates the ventral brainstem in the vicinity of the PreBoetz nucleus/caudal CO2 chemosensory area during middle age and dies again in old age. Ablating Cx26 genetically specifically within this population of about 20 cells by two independent neural crest Cre-driver lines leads to a loss of local ATP release in the posterior chemosensory area as well as a 40% reduction in elevated tidal volume responses specifically in middle age, as measured by whole-body plethysmography. These *in vivo effects* change over a life-time: they directly mirror the arrival, wiring in middle age and later death of this cell population in old age. This is a first known example of a middle age change in cranial neural crest lineage composition and highlights the significant power of very few cells for global metabolism. Such lineage-dependent middle-age dynamics also impacts upon the evolution of eusociality: altricial naked pups of our common amniote/synapsid ancestors were heated by by the breath of their (middle-aged) carers, sensing elevated CO_2_ levels in hypercapnic burrows. Such mechanistic exaptation enabled small amniotes to sense and survive the lethal global CO_2_ spikes during the Permo-Triassic and other extinction events.

## Introduction

Breathing is a vital function that maintains the partial pressures of O_2_ and CO_2_ in the arterial blood within the physiological limits. Chemosensory reflexes regulate the frequency and depth of breathing to ensure homeostatic control of blood gases. Historically, the ventral surface of the medulla oblongata has been recognized as a location of important central respiratory chemosensors (Loeschcke, 1982; Mitchell et al., 1963; Schlaefke et al., 1979; Schlaefke et al., 1970; Trouth et al., 1973). Recent work has focused on the retrotrapezoid nucleus (RTN) where a population of neurons highly sensitive to changes in pH/*P*CO_2_ has been described (Guyenet et al., 2008; Kumar et al., 2015; Mulkey et al., 2006; Mulkey et al., 2004) and the medullary raphé neurons, which appear also to be involved in detecting respiratory acidosis (Brust et al., 2014; Ray et al., 2011; Richerson, 2004). ATP-mediated signaling also contributes to the detection of pH/CO_2_. In response to hypercapnia, chemosensitive cells at the ventral surface of the medulla release ATP (Gourine et al., 2010; Gourine et al., 2005; Huckstepp et al., 2010b; Mulkey and Wenker, 2011; Wenker et al., 2010). This triggers adaptive ventilatory responses by exciting neurons of the brainstem respiratory network via both P2X and P2Y receptors (Gourine et al., 2003; Lorier et al., 2007).

According to traditional consensus, CO_2_ is detected via the consequent change in pH, and pH is a sufficient stimulus for all adaptive changes in breathing in response to hypercapnia (Loeschcke, 1982). pH-sensitive K^+^ channels (TASKs and KIRs) are potential transducers. Although TASK-1 in the peripheral chemosensors of the carotid body (CB) contributes to overall pH/CO_2_ chemosensitivity (Trapp et al., 2008), TASK-1 does not appear to play a role in central pH/CO_2_ chemosensing (Mulkey et al., 2007). By contrast TASK-2 may act as a central sensor of pH and contribute to adaptive changes in breathing (Kumar et al., 2015; Wang et al., 2013). Recently a pH sensitive receptor, GPR4, has been linked to central chemosensitivity in the RTN. Complete deletion of this gene (from all tissues) greatly reduces the CO_2_ chemosensitivity in mice (Kumar et al., 2015). However the effect of deletion of GPR4 is manifest mainly as an effect on the adaptive changes in respiratory frequency with only weak effects on CO_2_-induced changes in tidal volume. This is surprising as chemosensory stimulation of central chemosensory areas including the RTN predominantly evokes adaptive changes in the tidal volume of breathing (Li et al., 1999; Marina et al., 2010). This suggests that other mechanisms of chemosensory transduction in addition to GPR4 and pH sensing may also be involved. Although current evidence supports the hypothesis that CO_2_ detection can occur via the proxy of pH, several other strands of published data suggest that CO_2_ can have additional independent effects from pH on central respiratory chemosensors (Eldridge et al., 1985; Huckstepp and Dale, 2011; Shams, 1985).

We recently showed that CO_2_ directly binds to connexin26 (Cx26) hemichannels and causes them to open. We have identified the critical amino acid residues that are necessary and sufficient for this process (Huckstepp et al., 2010a; Meigh et al., 2013). This direct gating of Cx26 is an important new mechanism underlying CO_2_-dependent ATP release (Huckstepp et al., 2010a) (Huckstepp et al., 2010b; Wenker et al., 2012) and provides a potential mechanism for the direct action of CO_2_ on breathing. We also recently demonstrated that a mutation in Cx26 (A88V) that underlies keratitis ichthyosis and deafness syndrome completely removes CO_2_ sensitivity from Cx26 and is associated with prolonged bouts of central apnea in human infants (Meigh et al., 2014) and have shown in subsequent misexpression approaches that medullary cells can acquire new ATP release characteristics and in dominant negative approaches, virus infected cells are compromised in this function (Brotherton et al, 2024). This still leaves the question open, what the exact cell type and lineage origins of these Cx26-CO2 sensing cells are. In a parallel strand of work examining the fate of neural crest cells in postnatal times utilizing well-established Cre-drivers for neural crest cell fate we had discovered a novel very small group of cells that are visible at 1 and 4 months on the surface of the posterior medulla. This is territory that is previously known to be devoid of any neural crest cells, entirely mesodermal. If one could show that these newly found cells express connexin 26, one can test their function by a neural-crest specific ablation of the floxed Cx26 transgene. In this study we utilize targeted deletion of the Cx26 gene to determine a direct causal link between Cx26 gating and the CO_2_-dependent regulation of breathing in mice. Cx26 homozygous mutants are early embryonic lethal, necessitating a conditional mutagenesis approach (Cohen-Salmon et al., 2002; Gabriel et al., 1998). We have ablated Cx26 in 4 different, genetically well defined cell populations. Our deletion of Cx26 provides the first genetic evidence that direct detection of CO_2_ by Cx26 is a major regulator of breathing *in vivo* in adult rodents at modest levels of hypercapnia. We have identified a unique (transient) class of Cx26+ chemosensitive cells derived from the neural crest (NCC), which are essential to the increased central CO_2_ chemosensitivity in middle age.

## Results

### Genetic excision of Cx26 in GFAP+ cells greatly reduces CO_2_-dependent ATP release at the ventral medullary surface and reduces respiratory CO_2_ chemosensitivity

Prior evidence suggested that GFAP+ cells in the brainstem express Cx26 (Huckstepp et al., 2010b). One major class of GFAP+ cell, astrocytes, has been implicated in central chemosensory responses (Gourine et al., 2010; Mulkey and Wenker, 2011; Wenker et al., 2010). We therefore carefully examined the colocalization of GFAP- and Cx26-immunoreactivity (IR) at and near the ventral surface of the medulla oblongata in both the rostral and caudal chemosensory areas as defined by Loeschke (Figure 1A). In the rostral area, Cx26-IR was extensively distributed in both the parenchyma, where it was associated with GFAP-IR, and in the leptomeninges (Figure 1B, Figure 1 Supplement 1). Non-chemosensory areas such as the pyramidal tracts do not harbour Cx26-IR (Figure 1 Supplement 1). Furthermore Cx26-IR was absent from the carotid body (Figure 1 Supplement 2). In GFAP-Cre:Cx26^fl/fl^ mutants, the Cx26-IR was abolished in the parenchyma but remained in leptomeninges, vasculature and arachnoid layers (Figure 1C, Figure 1 Supplement 3). This confirms the specificity of the Cx26 antibody, the specific localization of Cx26 in GFAP+ cells and the efficacy of GFAP-Cre mediated Cx26 ablation (Garcia et al., 2004).

**Figure 1.**
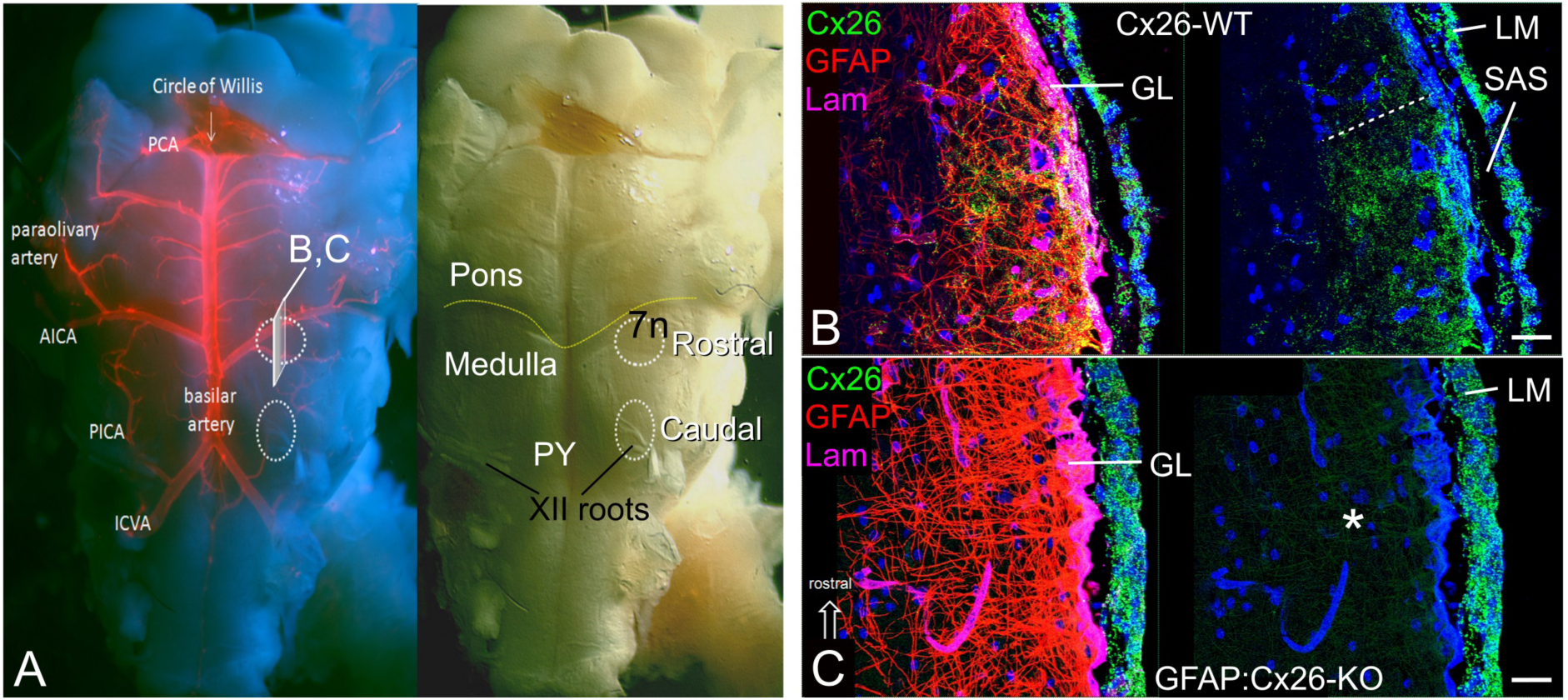
Cx26 expressed by GFAP+ cells at the ventral medullary surface. A) Wholemount picture of the medulla oblongata of mouse showing major blood vessels stained with lectin (left) and position of the major external landmarks along with the rostral and caudal chemosensing areas (right). B) Parasagittal section of medulla from wild type mouse (position and plane indicated in (A)) showing GFAP (red), laminin (magenta) and Cx26 (green) immunoreactivity, reaching 87 ± 10 µm into the rostral WT chemosensitive area (dotted line). GL –glia limitans, LM –leptomeninges, SAS –subarachnoid space. Image on the right comprises only Hoechst staining (blue) plus Cx26, to show the widespread distribution of Cx26 more clearly. C) Equivalent parasagittal section from GFAP:Cx26-KO mouse, rostral chemosensitive area. Cx26 immunoreactivity remains in the leptomeninges, but is completely absent from parenchyma (asterisk) demonstrating that it is normally expressed in GFAP+ cells. Scale bars in (B and C) 20 µm.

We tested the effect of ablating Cx26 from GFAP+ cells on CO_2_-evoked ATP release in the rostral and caudal chemosensory areas of the ventral medullary surface, as ATP may act as a critical mediator of central respiratory CO_2_ chemosensitivity (Gourine et al., 2005). CO_2_-induced ATP release was significantly reduced in the rostral (p<0.001) and caudal chemosensory (p<0.05) areas of brainstem horizontal slices derived from 19 GFAP-Cre^+/-^:Cx26^fl/fl^ (GFAP:Cx26-KO) mice as compared to 20 of their GFAP-Cre^-/-^:Cx26^fl/fl^ (GFAP:Cx26-WT) litter mates that were wild-type for Cx26 (Figure 2A,B, Table 1). This shows that Cx26 is a major conduit for CO_2_-evoked ATP release in the medulla oblongata. The remaining leptomeningeal Cx26 in these mutants does not appear to contribute markedly to the recorded CO_2_-evoked ATP release in our preparations.

**Figure 2.**
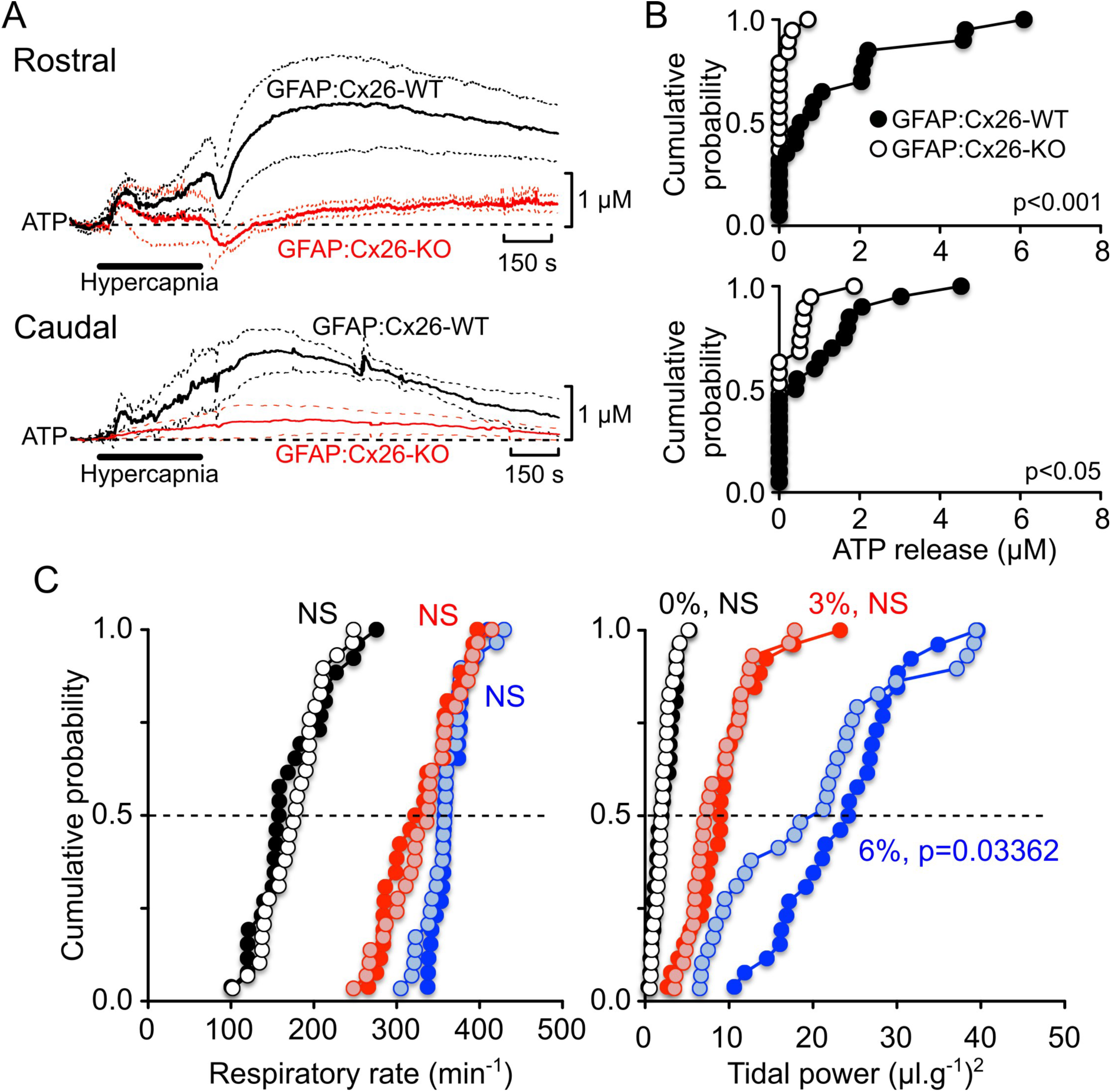
Cx26 in GFAP+ cells contributes to the chemosensory control of breathing. A) Comparison of CO_2_-evoked ATP release recorded with biosensors from horizontal slices of ventral medullary surface from GFAP:Cx26-WT and GFAP:Cx26-KO mice. The selective deletion of Cx26 from GFAP+ cells greatly reduces CO_2_-evoked ATP release from both the rostral (top) and caudal (bottom) chemosensory areas. Each trace is an average of 5 experiments, dashed line shows the standard error. B) Cumulative probability distributions show that in the rostral area (top), deletion of Cx26 has completely abolished ATP release, whereas in the caudal area (bottom) a small amount of ATP release remains. C) Cumulative probability plots of respiratory rate and tidal power for GFAP:Cx26-WT and GFAP:Cx26-KO mice at 0 (black circles), 3 (red circles) and 6% (blue circles) inspired CO_2_ – the lighter filled circles represent the knockout mice at each level of CO_2_. There is no difference in respiratory rate between wild type and knockout mice at any level of inspired CO_2_. The tidal power of breathing for the knockout mice is significantly less than the wildtype mice at 6% inspired CO_2_, but no different at any other level of inspired CO_2_.

**Table 1.** Median CO_2_-dependent ATP release determined for GFAP:Cx26-WT (GFAP-Cre^-/-^:Cx26^fl/fl^) and GFAP:Cx26-KO (GFAP-Cre^+/-^:Cx26^fl/fl^) littermates and Wnt1:Cx26-WT (Wnt1-Cre^-/-^:Cx26^fl/fl^) and Wnt1:Cx26-KO (Wnt1-Cre^+/-^:Cx26^fl/fl^) littermates. The medians are shown with the 95% upper and lower confidence limits in parentheses underneath. (* p<0.05, **p<0.01, *** p<0.001, Mann Whitney U test between wild type and knock outs).

| Genotype | Rostral | Caudal | n |
| --- | --- | --- | --- |
| GFAP:Cx26-WT | 0.5***<br>(0, 2.1) | 0.4*<br>(0, 1.3) | 20 |
| **GFAP:Cx26-KO** | **0\*\*\***<br>**(0, 0)** | **0\***<br>**(0, 0.56)** | 19 |
| Wnt1:Cx26-WT | 0.2<br>(0, 0.6) | 0.7**<br>(0.2, 1.7) | 17 |
| **Wnt1:Cx26-KO** | **0**<br>**(0, 0.3)** | **0\*\***<br>**(0, 0.3)** | 20 |

We next used whole body plethysmography (WBP) in conscious unrestrained mice to record CO_2_-induced responses *in vivo* in 29 GFAP:Cx26-KO and 26 GFAP:Cx26-WT mice derived from 4 litters. At all levels of applied inspired CO_2_, there was no significant difference in the respiratory frequency between the GFAP:Cx26-KO and their GFAP:Cx26-WT littermates: both genotypes exhibited identical increases in the respiratory frequency in response to increases in the inspired CO_2_ (Figure 2C, Table 2).

**Table 2.** Median respiratory rate and tidal power determined by whole body plethysmography for: GFAP:Cx26-WT (GFAP-Cre^-/-^:Cx26^fl/fl^) and GFAP:Cx26-KO (GFAP-Cre^+/-^:Cx26^fl/fl^) littermates; Wnt1:Cx26-WT (Wnt1-Cre^-/-^:Cx26^fl/fl^) and Wnt1:Cx26-KO (Wnt1-Cre^+/-^:Cx26^fl/fl^) littermates; P0:Cx26-WT (P0-Cre^-/-^:Cx26^fl/fl^) and P0:Cx26-KO (P0-Cre^+/-^:Cx26^fl/fl^) littermates; and PGDS:Cx26-WT (PGDS-Cre^-/-^:Cx26^fl/fl^) and PGDS:Cx26-KO (PGDS-Cre^+/-^:Cx26^fl/fl^) littermates. The medians are shown with the 95% upper and lower confidence limits in parentheses underneath. At 6% inspired CO_2_ all the GFAP:Cx26-KO, Wnt1:Cx26-KO and P0:Cx26-KO knock out strains of mice show significantly less tidal power than the wild types (* p<0.05, *** p<0.001, Mann Whitney U test between wild type and knock outs). Posthoc statistical power achieved at p=0.05 for GFAP-Cre comparison at 6% CO_2_, 86%; for Wnt1-Cre comparison at 6% CO_2_ 96%; and for P0-cre comparison, were effect size to be same as Wnt1-Cre at 6%, 87%.

| %CO<sub>2</sub> | Respiratory rate (min<sup>-1</sup>) |  |  | Tidal power (μl.g<sup>-1</sup>)<sup>2</sup> |  |  | n |
| --- | --- | --- | --- | --- | --- | --- | --- |
|  | 0 | 3 | 6 | 0 | 3 | 6 |  |
| GFAP:Cx26-WT | 158<br>(150, 206) | 322<br>(286, 357) | 358<br>(355, 376) | 2<br>(1.1, 3.1) | 9<br>(7.2, 10.7) | 24.2*<br>(19.1, 27.6) | 26 |
| **GFAP:Cx26-KO** | **178**<br>**(157,194)** | **338**<br>**(311, 358)** | **358**<br>**(348, 373)** | **1.9**<br>**(1.5, 2.6)** | **7.4**<br>**(6, 10.8)** | **18.1\***<br>**(10.9, 24.1)** | **29** |
| Wnt1:Cx26-WT | 195<br>(175, 211) | 302<br>(283, 317) | 347<br>(336, 355) | 1.8<br>(1.2, 2.0) | 5.1<br>(3.9, 6.5) | 17.6***<br>(14.6, 24.6) | 48 |
| **Wnt1:Cx26-KO** | **197**<br>**(190, 217)** | **308**<br>**(293, 327)** | **335**<br>**(324, 353)** | **1.4**<br>**(1.1, 1.6)** | **4.6**<br>**(2.7, 6.1)** | **10.2\*\*\***<br>**(7.1, 16.6)** | **46** |
| P0:Cx26-WT | 183<br>(119, 229) | 357<br>(285, 391) | 357<br>(335, 372) | 1.5<br>(0.65, 1.9) | 9.2<br>(5.5, 18.3) | 28.5*<br>(22.4, 43.1) | 8 |
| **P0:Cx26-KO** | **175**<br>**(137, 243)** | **356**<br>**(322, 392)** | **358**<br>**(339, 378)** | **1.4**<br>**(0.7, 2.2)** | **8.9**<br>**(5.6, 11.8)** | **20.9\***<br>**(16.7, 30.4)** | **11** |
| PGDS:Cx26-WT | 178<br>(158, 193) | 326<br>(304, 350) | 338<br>(322, 341) | 1.2<br>(0.93, 1.4) | 7.5<br>(4.9, 9.7) | 21.2<br>(14.4, 27.5) | 30 |
| **PGDS:Cx26-KO** | **209**<br>**(165, 226)** | **318**<br>**(284, 339)** | **338**<br>**(320, 342)** | **1.3**<br>**(0.99, 1.9)** | **8.8**<br>**(6.1, 11.6)** | **19.8**<br>**(13.1, 26.7)** | **30** |

Analysis of tidal power at the eupneic breathing frequency (a measure related to eupneic tidal volume, but obtained in the frequency- rather than the time domain) showed that there was no significant differences between the two genotypes at 0% and 3% inspired CO_2_. However at 6% inspired CO_2_, the GFAP:Cx26-KO mice exhibited significantly lower tidal power than the GFAP:Cx26-WT mice (Figure 2C, Table 2). Interestingly, the differences between the GFAP:Cx26-KO and GFAP:Cx26-WT mice were most obvious around the lower quartile of the cumulative probability distribution and much less apparent around the upper quartile. This suggests phenotypic variability. We do not know the origins of this variability, but it might arise from varying degrees of compensation. As will be seen below, a more restricted genetic ablation of Cx26 yields a more uniform WBP phenotype.

### A novel GFAP+ neural crest cell type on the ventral surface of the caudal chemosensitive area of the medulla oblongata

To home in more closely on the specific cell type within the larger GFAP+ cell population that is Cx26+, responsible for ATP release and the WBP mutant phenotype we observed in the GFAP conditional Cx26 mutants, we first characterized the cellular origins of the chemosensory regions with regard to expression of Cx26. While Cx26-IR in the rostral region is widespread among the GFAP+ cells (Figure 1B), only very few cells of the caudal region display Cx26-IR inside the brain parenchyma (Figure 3A,B). The developmental origins and anatomy of these caudal cells remained unclear.

**Figure 3.**
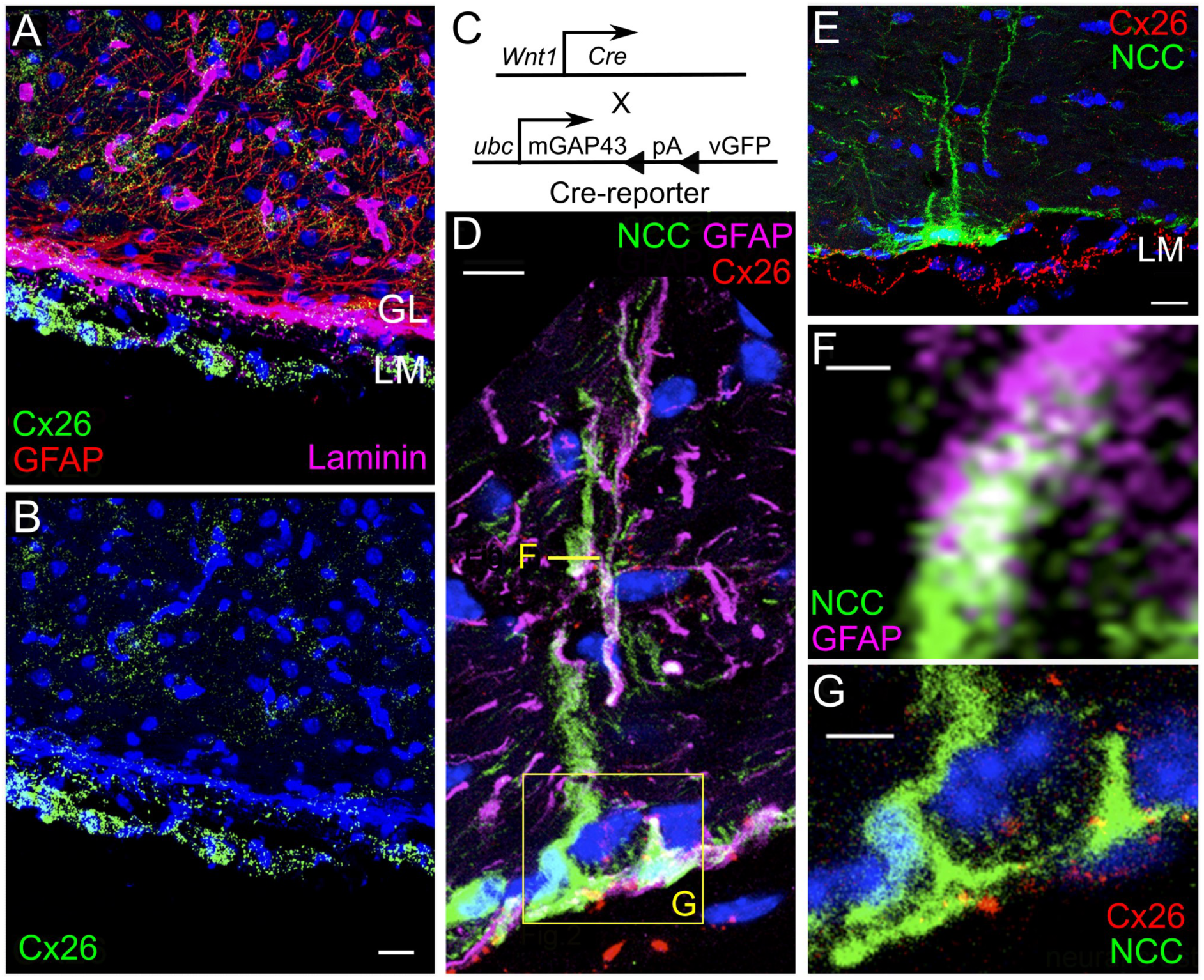
In the caudal CO_2_ chemosensory area, neural crest derived cells express Cx26. A and B) Single optical section showing scarce intraparenchymal and abundant leptomeningeal (LM) Cx26-IR (green), GFAP-IR in red and laminin-IR (magenta) labelling the ‘glia limitans’ (GL). C) Genetic lineage labelling strategy to reveal neural crest cell anatomy (Matsuoka et al., 2005), using a new recombinase reporter (see Materials and Methods). D-G) Anatomical characterization of the GFAP+ NCCs in caudal medulla. D and E) NCCs genetically marked (vGFP, green) reach deep into the brain parenchyma and express Cx26 (red) and GFAP (magenta), confocal z-stacks. F,G) Single optical sections showing that the same NCCs express both GFAP and Cx26. Scale bars: A, B) 20 µm; D, 10 µm; E, 20 µm; F, 2 µm; and G, 10 µm.

We reasoned that neural crest cells (NCCs) might express Cx26 and test this using the well-established Wnt1-Cre- and P0-Cre- transgenes that are specific for NCCs (Danielian et al., 1998; Matsuoka et al., 2005; Yamauchi et al., 1999) (Figure 1 Supplement 3). Although NCCs are currently thought to be leptomeningeal in the fore and midbrain regions, but not considered to be present in this particular medullary brain region (Etchevers et al., 2002; Etchevers et al., 2001), their distribution at adult stages has never been examined. Surprisingly, we found genetically tagged (GFP+) NCC to be present only in the caudal, but not in the rostral, medullary chemosensory area (Figure 3D,E).

We examined whether NCCs in the caudal region express Cx26. by utilizing a novel recombinase reporter transgene, which accomplishes membrane-localization of vGFP via a GAP43-membrane anchor (Figure 3C). This visualizes the entire anatomy of the permanently marked NCCs (Figure 3D-G), which can be reconstructed at high resolution in 3D (Figure 3D, F, G, Supplementary Movie 1), and clearly shows puncta of Cx26 protein on the surface of these cells (Figure 3G). While cellular somata are embedded in the glia limitans (GL) each cell sends a single process of 119 ± 6 µm into the brain parenchyma of the caudal medulla oblongata. The genetic labeling of the entire cellular surface with membrane-bound GFP allowed us to observe that all of the Cx26+ NCCs express GFAP. Each cell had a single process, which displayed a spatulate morphology but was only partially filled with GFAP protein (Figure 3F). The processes of GFAP+ NCCs were found to be 3 times wider than those of other (astrocytic) GFAP+ cells localized deeper inside the brain parenchyma. We could only find a maximum of 25 Cx26+/GFAP+ NCCs per caudal brain region per animal (n=5 animals). These neural crest cells were restricted to a small area just anterior to the first, most anterior hypoglossal (XIIn) root, directly overlying the pre-Bӧtzinger complex, a key respiratory rhythm generator. This tight localization of Cx26+ NCCs is strikingly similar to the chemosensitive Schlaefke (S) area (Schlaefke et al., 1970). Apart from the leptomeninges, the only cells that we identified as Cx26+ in this caudal region were NCCs (Table 3). As described below, fore- and midbrain leptomeningeal cells are Cx26-IR, but selective ablation of the gene from these cells had no discernable *in vivo* effect.

**Table 3.** Localization of Cx26 in medullary cell types and structures. The assessment criteria were colocalization of Cx26 immunoreactivity (Cx26-IR) with genetically tagged cells and loss of Cx26 immunoreactivity after cell specific gene ablation. The Wnt1-Cre, P0-Cre and GFAP-Cre ablations have only one cell type in common that is also Cx26-IR (shaded in grey) and can thus yield the conditional mutant phenotype.

| Structures at 4 months | Embryonic origins |  |  |
| --- | --- | --- | --- |
|  | Mesoderm | Neural crest (genetically labeled with Wnt1-Cre and P0-Cre) | CNS |
| Medullary LM and subarachnoid space | Cx26-IR |  |  |
| Blood vessel endothelia | Cx26-IR |  |  |
| Superficial GFAP+ cells (caudal area only) |  | Cx26-IR, Genetically labelled with GFAP-Cre |  |
| Astrocytes of rostral area |  |  | Cx26-IR, Genetically labelled with GFAP-Cre |
| Astrocytes of caudal area |  |  | Genetically labelled with GFAP-Cre |
| Carotid body |  | Genetically labelled with GFAP-Cre |  |

Thus any *in vivo* effects that could be seen from a neural-crest specific gene ablation would have to be assigned to the intersection of neural crest origin in the medulla and Cx26 gene expression.

### NCC-specific Cx26 ablation specifically reduces tidal volume increases of breathing during hypercapnia

The genetic identification of these unique Cx26+ NCCs allowed us to selectively ablate Cx26 from them by crossing the Cx26^fl/fl^ strain with the same Wnt1-Cre driver and vGFP reporter transgenes (Figure 4A). In these crosses, only cells that are vGFP-labelled are thus expected to also have lost the Cx26 gene *in vivo*. In the resultant homozygous Cx26 mutants we still observed vGFP+ NCCs, indicating that their ontogeny was not affected by ablation of Cx26 and is therefore Cx26-independent. Cx26-IR was specifically lost from these (GFP+) NCCs in Wnt1-Cre^+/-^:Cx26^fl/fl^ mice (Wnt1:Cx26-KO), while leptomengineal Cx26 expression remained, demonstrating high cell-selectivity of this genetic ablation approach (Figure 4B-D). This also confirms that leptomeninges surrounding the medulla are not neural crest derived, thus could not contribute towards any *in vivo* phenotype.

**Figure 4.**
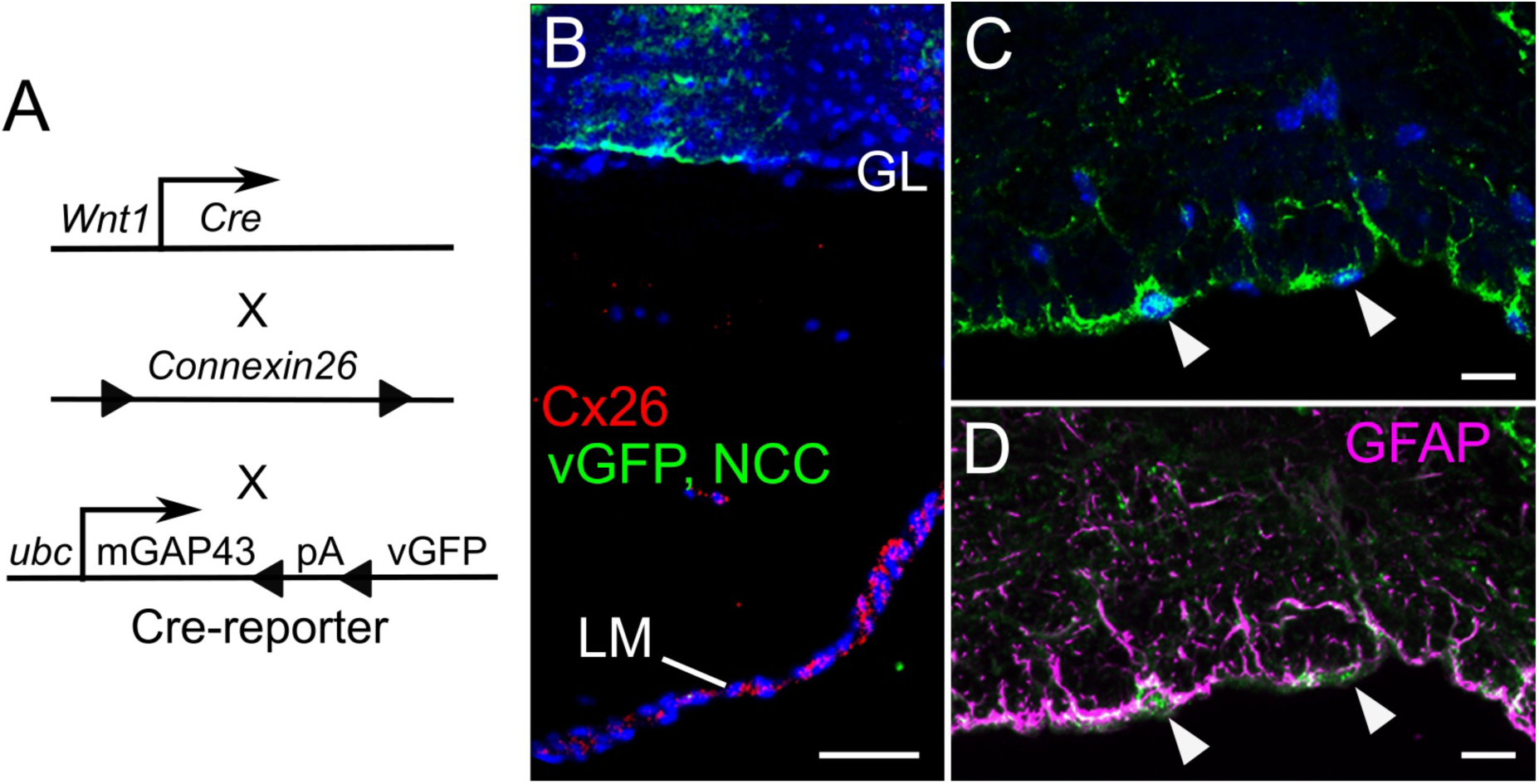
Deletion of Cx26 from the neural crest cells in the caudal chemosensory area. A) Genetic scheme to effect neural crest selective knockout of Cx26 and simultaneous marking of the neural crest derived cells shown in (B-D). B) Absence of Cx26 from the GFP+ NCCs (marked with vGFP, green), but its continued presence in the (non-neural crest, vGFP-) leptomeninges (LM). C, D) GFP+ NCCs cells do not express Cx26 but are GFAP+ (magenta). Arrowheads in (C, D) indicate the cell bodies of marked neural crest cells at the ventral surface of the medulla in different channels (green vGFP). Scale bars B) 100 µm, C, D) 20 µm.

Having established successful deletion of Cx26 from the small number of NCCs in the caudal chemosensory area, one would expect altered ATP release in this area due to the CO_2_-dependent gating of Cx26 that we previously established. We tested the effect on focal CO_2_ –evoked ATP release in this very restricted (caudal) area using acute horizontal slices of the ventral medullary surface. Indeed, ATP release in response to hypercapnia was reduced in the caudal chemosensitive area (slices from 20 Wnt1:Cx26-KO and 17 Wnt1-Cre^-/-^:Cx26^fl/fl^ (Wnt1:Cx26-WT) mice), but was unaffected in the rostral chemosensitive area (which does not contain Cx26 expressing NCCs, Figure 5A,B, Table 1). In the caudal area the Wnt1-Cre driven excision of Cx26 yielded a similar phenotype compared to that of the GFAP-Cre driven Cx26 deletion (compare Figures 5A and 2A). This is consistent with the observation that all Cx26 expressing NCCs are also GFAP+. While a GFAP-Cre mediated gene ablation could also potentially affect GFAP+ astrocytes, Wnt1-Cre neural crest mediated ablation cannot, because the transgene is not expressed in astrocytes or their progenitors.

**Figure 5.**
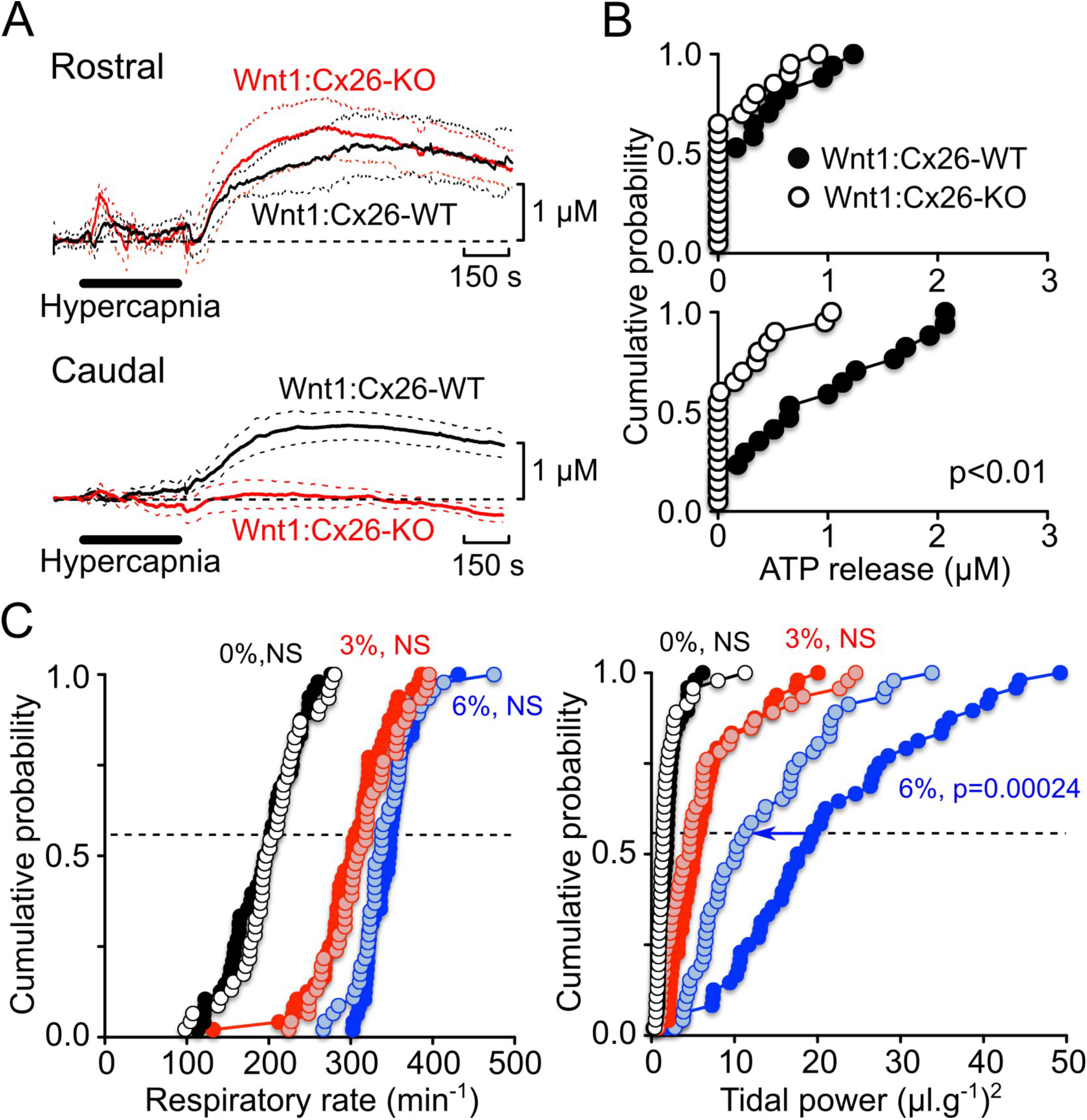
Cx26 in neural crest cells in the caudal medulla contributes to the chemosensory control of breathing. A,B) Biosensor recordings from horizontal slices of the ventral medullary surface show that knockout of Cx26 from neural crest cells has no effect on CO_2_-evoked ATP release in the rostral chemosensory area, but greatly reduces it from the caudal chemosensory area. Each trace is an average of 5 experiments, dashed line shows the standard error. c) Cumulative probability plots of respiratory rate and tidal power for Wnt1:Cx26-WT and Wnt1:Cx26-KO mice at 0 (black circles), 3 (red circles) and 6% (blue circles) inspired CO_2_ –the lighter filled circles represent the knockout mice at each level of CO_2_. There is no difference in respiratory rate between wild type and knockout mice at any level of inspired CO_2_. The tidal power of breathing for the knockout mice is significantly less than the wild type mice at 6% inspired CO_2_, but no different at any other level of inspired CO_2_.

To examine the *in vivo* effect of this focal Cx26 ablation inside NCCs of the caudal medulla we performed WBP on 48 Wnt1:Cx26-KO and 46 Wnt1:Cx26-WT mice from 7 different litters. There was no difference in respiratory frequency at the different levels of inspired CO_2_ (0, 3 and 6%) between Wnt1:Cx26-KO and Wnt1:Cx26-WT (Figure 5C). However, the increases in the depth of breathing (tidal volume, plotted as tidal power) in response to 6% hypercapnia were greatly reduced in Wnt1:Cx26-KO mice compared to Wnt1:Cx26-WT mice (p=0.0002, Figure 5C, Table 2). Overall the median tidal power at 6% inspired CO_2_ was reduced by 42% in the Wnt1:Cx26-KO mice compared to the Wnt1:Cx26-WT mice. We also controlled for potential genetic background effects linked to the Wnt1-Cre allele by performing WBP under WT Cx26 conditions. The presence of the Wnt1-Cre allele by itself on a WT Cx26 background did not alter ventilatory responses to hypercapnia (Figure 5 Supplement 1), demonstrating that it was the deletion of Cx26 rather than the presence of the Cre recombinase that gave rise to the phenotype.

The remarkable restriction of Cx26 to a caudal NCC population and the resulting ATP-release and WBP phenotypes in the NCC specific mutants called for an independent confirmation of the *in vivo* phenotype. To this end we utilized a second independent NCC-specific Cre-recombinase driver allele, P0-Cre. The P0-Cre transgene marks post-migratory NCCs only after they have left the neural tube and does not show expression inside the CNS. Thus although the total numbers of NCC labeled with this transgene are lower than that of the Wnt1-Cre transgene, direct CNS effects can be fully excluded and any similarity in phenotype to the Wnt1-Cre mediated Cx26 ablation has to be attributed to the newly discovered neural crest cell type. We compared responses to inspired CO_2_ in 11 P0-Cre^+/-^:Cx26^fl/fl^ (P0:Cx26-KO) mice and 8 P0-Cre^-/-^:Cx26^fl/fl^ (P0:Cx26-WT) littermates. Similar to the Wnt1:Cx26-KO mutants, the P0:Cx26-KO mutants exhibited changes in respiratory frequency in response to increased inspired CO_2_ at 6% that were no different from those of the P0:Cx26-WT mice. The tidal power at 0% and 3% were also similar between the mutant and wild type mice. However, the P0:Cx26-KO mice exhibited a tidal power that was significantly less than that of the P0:Cx26-WT littermates (p<0.05, 27% reduction of median tidal power, Figure 5 Supplement 2, Table 2). The similarity in phenotype between Wnt1-Cre and P0-Cre mediated Cx26 ablations consolidates the physiological significance of the newly discovered chemosensory NCCs in the ventilatory response to CO_2_ levels at 6%.

As described above, Cx26 is expressed in leptomeninges of the fore- and midbrain and as these are neural crest derived it is possible in principle that this leptomeningeal expression could affect in some ways the global respiratory response via mechanisms beyond the medulla oblongata. (Figure 1 Supplement 2). To assess a potential contributing factor of Cx26+ leptomeningeal cells towards CO_2_ chemosensing we sought to delete Cx26 globally in all leptomeningeal cells. We therefore crossed the PGDS-Cre mouse line to the floxed Cx26 mouse line. The PGDS-Cre transgene is expressed in all leptomeningeal cells, fully labels this tissue genetically and is not expressed inside the brain (Kalamarides et al., 2011). Thus a PGDS-Cre:Cx26^fl/fl^ cross deletes Cx26 from the entire leptomeningeal layer without affecting any other cell types in the brain (Kalamarides et al., 2011) (Figure 1 Supplement 2). We compared responses to inspired CO_2_ in PGDS-Cre^+/-^:Cx26^fl/fl^ (PGDS:Cx26-KO) mice and PGDS-Cre^-/-^:Cx26^fl/fl^ (PGDS:Cx26-WT) littermates. This leptomeningeal-specific deletion of Cx26 had no effects on the chemosensitivity of breathing to inspired CO_2_ (Figure 5 Supplement 3, Table 2). We infer from this that despite being Cx26 positive the (mesodermal and neural crest derived) leptomeninges do not contribute additionally towards the respiratory response to hypercapnia via the actions of Cx26.

We infer from the Wnt1-Cre, P0-Cre, GFAP-Cre and PGDS-Cre mediated ablations of Cx26 that superficial GFAP+ NCCs, restricted to the caudal part of the ventral medullary surface, express Cx26 and are responsible for the CO_2_-dependent ATP release in this region. Genetic Cx26 ablations inside these cells allow us to link CO_2_-evoked Cx26–dependent ATP release in the caudal region to the central control of tidal volume in awake conscious animals.

### Postnatal ontogeny of Cx26-mediated respiratory CO_2_ chemosensing: a new focus on middle age

The functional experiments outlined so far only examined central respiratory chemosensitivity in 3-4 months old mice, customary for studies of adult chemosensing. Given the unusual origin and localization of these NCC chemosensors, we were curious to know if Cx26-mediated CO_2_ chemosensing changes with age. We therefore examined CO_2_- induced responses across the lifespan: in Wnt1:Cx26-KO and Wnt1:Cx26-WT mice at 2, 3, 4 and 9 months. Over this age progression the response to CO_2_ in terms of changes in respiratory rate remained very similar and there were no differences between the knockout and wild type littermates (Figure 6). However, tidal power responses were successively greater at 2, 3 and 4 months before decreasing again at 9 months (Figure 6A). A similar age-related change in CO_2_ sensitivity has been previously described in mice: at 32-44 weeks mice exhibit smaller hypercapnic responses than at the age of 10-16 weeks (Onodera et al., 1997). What is striking is that at 2 and 9 months there was no difference in the tidal power at 6% inspired CO_2_ between the wild type and knockout mice. Thus a Cx26 mediated (neural crest) mechanism might not act in those periods of the postnatal ontogeny/lifespan. However, at 3 and 4 months the tidal power at 6% inspired CO_2_ was significantly less in the Wnt1:Cx26-KO mice than in their Wnt1:Cx26-WT littermates (Figure 6A). We interpret this as the emergence of an additional direct CO_2_ sensing pathway at middle-age. To verify the anatomical location of these changes in chemosensitivity we tested whether ATP was released from the caudal area of ventral medullary slices derived from a similar range of ages of wild type mice (Figure 6B). In mice aged between 2 and 3 months, CO_2_-evoked ATP release was observed in 2/11 slices (median 0 µM, 95% confidence interval 0 to 1.2 µM). In slices from mice more than 7 months old, ATP release was seen in only 1/21 slices (median 0 µM, 95% confidence interval 0 to 0 µM). By contrast in 15 slices from mice aged between 3 and 7 months of age the median CO_2_-evoked ATP release was 1.1 µM (95% confidence interval 0.9 to 1.8 µM). This increased ATP release probability matches to the postnatal development of the WBP phenotypes, the increased chemosensitivity between 2 and 7 months in the awake animal.

**Figure 6.**
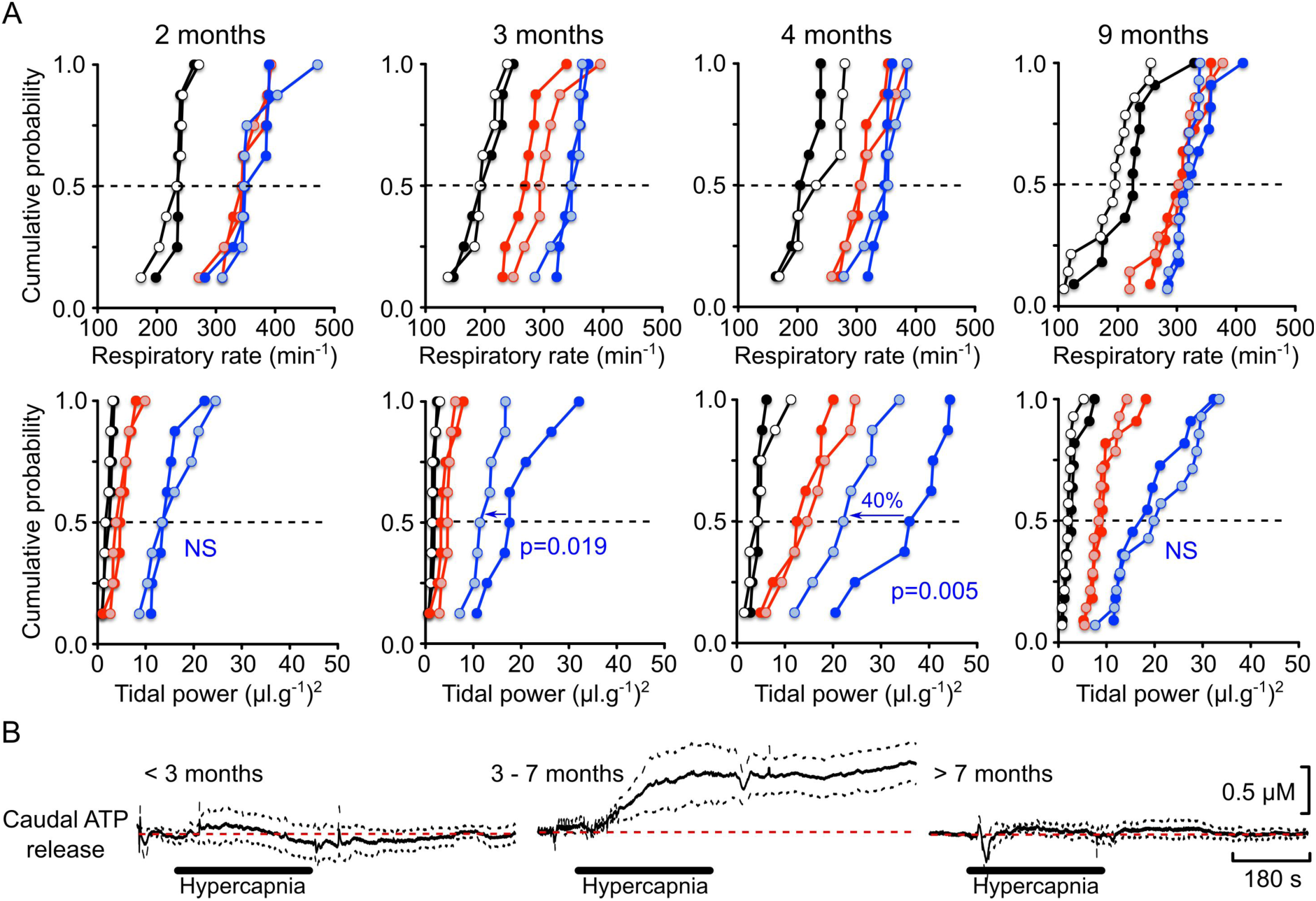
Postnatal ontogeny of the Cx26-dependent CO_2_ chemosensitivity: a role in middle age. A) Wnt1:Cx26-WT and Wnt1:Cx26-KO littermates were assessed at 2, 3, 4 and 9 months of age by WBP. Same litter for 2, 3 and 4 months, a different litter for 9 months. Data displayed as cumulative probability graphs for respiratory rate and tidal power (black circles 0%, red circles 3 %, blue circles 6%, the lighter colour indicating the knockout mice). Over this period altering inspired CO_2_ gave similar adaptive responses in respiratory rate (top row) between wild type and knock out littermates. However the adaptive response in tidal power at 6% inspired CO_2_ was affected (bottom row). At 2 months, no phenotype was evident with Wnt1:Cx26-KO. When tested at 3 and 4 months, compared to wild types, the Wnt1:Cx26-KO mutants exhibited less tidal power at 6% CO_2_ than the Wnt1:Cx26-WT. At 9 months, there was no effect of deletion of Cx26 on the respiratory response to 6% CO_2_. B) Averages (5 recordings) of ATP biosensor recordings showing lack of CO_2_ evoked (black bar) ATP release from ventral surface of medullary slice (cut from wild type mice) at ages less than 3 months or greater than 7 months. Robust ATP release was seen in slices cut from mice from 3 to 7 months of age.

### The presence of NCC CO_2_ chemosensors in the caudal medulla is restricted to middle age

Could the natural developmental increase in respiratory CO_2_ sensitivity be explained by an age-dependent presence of NCC-derived chemosensory cells, or is it related to epigenetically controlled differential Cx26 gene expression over time inside circuitry that is already present at birth? To discriminate between these possibilities, we genetically tracked NCC lineages over a corresponding time course and studied their Cx26 immunoreactivity. Wnt1-Cre:confetti reporter (Snippert et al., 2010) crosses permitted us to investigate clonal relationships (Figure 7A) in the ventral medulla between 1 and 8 months: cells belonging to the same clone carry the same genetically encoded fluorescent colour. At no period were we able to detect NCCs in the rostral chemosensory region. We detected the first NCCs in the caudal region at 1 month, residing at the surface of brain parenchyma, being Cx26+ and each cell carrying many processes (Figure 7B,C). At this early stage Cx26-IR is restricted in this area to NCCs (white arrows in Figure 7B) and adjacent (mesodermal) leptomeninges. Figure 7C shows clonal expansions at 1 month (RFP- versus GFP-tagged clones), but the Cx26+ NCCs do not yet display their adult morphology with each cell carrying a single radial spatulate process. This only happens after 3 months (Figure 7D) when each cell carries a single process reaching into the brain parenchyma, thus a considerable time after attainment of sexual maturity (8 weeks in the C57/B6 strain used). To our surprise the cells were no longer detectable after 8 months: only in rare instances did a few flat NCCs remain (without radial processes) in the caudal region (Figure 7E). Thus the emergence (at 3 months) and the loss (after 8 months) of the Cx26-mediated hypercapnic tidal volume response in awake animals closely matches the appearance, maturation in middle-age, and subsequent loss of the superficial Cx26+ neural crest cells that we identified in the caudal chemosensory region of the medulla oblongata (summarized in Figure 8).

**Figure 7.**
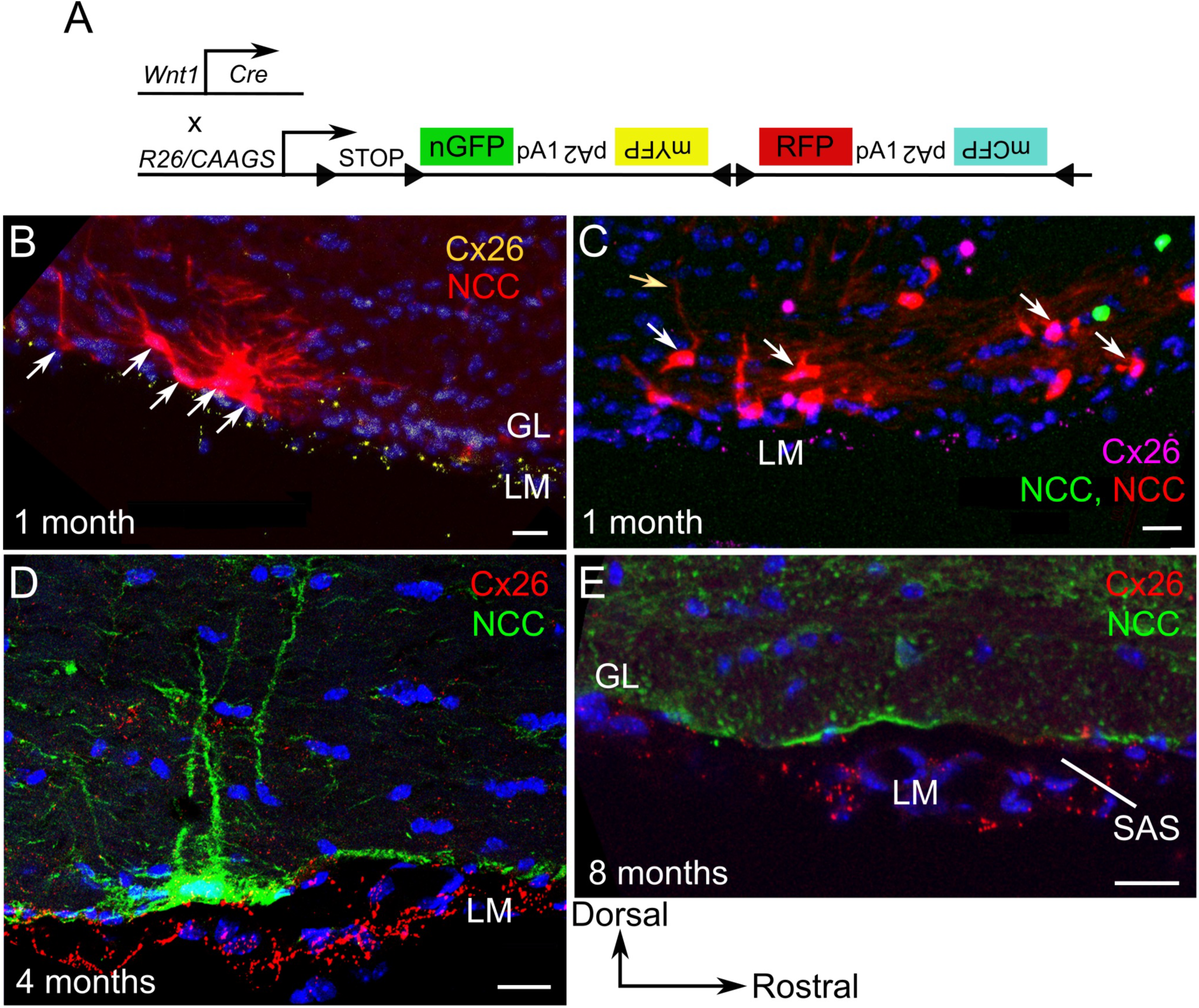
Arrival of the neural crest chemosensory cells in the caudal medulla parallels the onset of functional Cx26-dependent CO_2_ chemosensitivity. A) Wnt1-Cre was crossed into the confetti reporter for clonal resolution. Clones (derived from same NC mother cell) express the same genetically encoded colour. B-E) Parasagittal sections in of the caudal chemosensitive area at 1, 4 and 8 months. B) At 1 month large NCCs (white arrows) display a complex tangential immature morphology and very low levels of Cx26-IR. C) Tangential section through caudal surface area: clonal expansion is discernable (a red clone (white arrows), and a green clone), low-level Cx26-IR on cellular soma, some cells develop single thin radial process (yellow arrow). D) At 4 months, they have developed a single fat radial process each and express Cx26 superficially and on their process. E) At 8 months, the neural crest cells have largely disappeared, leaving only labeled membrane fragments. The few remaining NCCs have lost their radial process. Cx26 expression is confined to the leptomeninges (LM). D and E) Labeled NCCs in Wnt1-*Cre* ubc-GAP43-vGFP^fl/fl^ mice. All scale bars 20 µm. SAS, subarachnoid space; GL, glia limitans.

**Figure 8.**
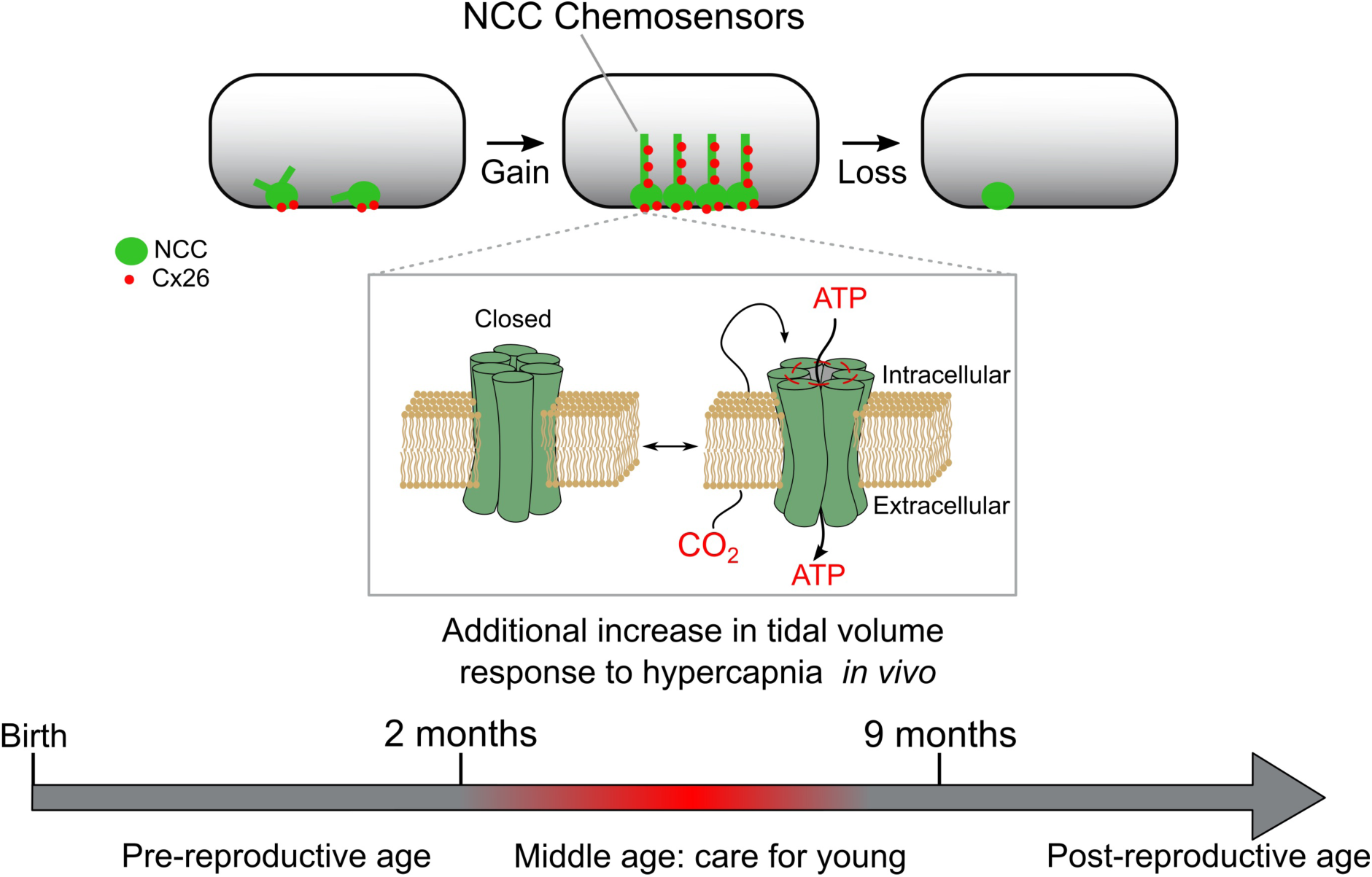
Conceptual diagram indicating proposed role of NCC Cx26+ chemosensors in respiratory chemosensitivity. The caudal area of the medulla oblongata indicated by grey shaded rectangle at 1,4 and 9 months. Early on, NCCs destined to mature into chemosensors (green) are present with some Cx26-IR (red circles) but lacking mature morphology. Post-puberty in middle age at peak of reproductive activity, the NCCs have mature morphology and strong expression of Cx26. CO_2_ binds directly to Cx26 (Meigh et al., 2013) to trigger hemichannel opening and release of ATP which gives enhanced adaptive respiratory responses to hypercapnia through bigger increases in tidal volume. Later on both the NCCs and Cx26-IR disappear from this area as the animal moves into a post-reproductive age, and respiratory chemosensitivity lessens.

## Discussion

### Genetic targeting strategy and rationale: xeroing in on the key cell population

We utilized four previously well-characterized Cre drivers to ablate Cx26 expression in the key cell populations involved in the phenotype. A GFAP-Cre transgene labels astrocytes and – as we show here - also a rare Cx26+ NCC population. Two independent neural crest Cre-drivers, the well-known Wnt-1Cre transgene labelling all premigratory neural crest (except that of rhombomere 1) (Matsuoka et al., 2005) as well as the P0-Cre driver labelling post-migratory neural crest (outside the CNS) (Yamauchi et al., 1999) yielded the same *in vivo* phenotype. A fourth driver, a PGDS-Cre transgene that ablates Cx26 in all leptomeninges (Kalamarides et al., 2011), irrespective of their neural crest or mesodermal origins (Figure 1 Supplement 2), gave no breathing phenotype.

Beyond the medulla oblongata, several other anatomical structures have been implicated in control of breathing or are neural crest in origin. However, any emergent Cx26 mutant phenotype can only be attributed to cell populations in which (or in whose mother cells) a Cre-driver is active and that also express Cx26 protein at the time of the physiological assays. This means that cells that are (or whose progenitors were) Cre positive but do not express Cx26 cannot contribute to a phenotype observed. As a case in point, we extensively queried whether the carotid body, a well-known regulator of breathing and a neural crest derivative, harbours Cx26+ positive cells. Despite our strenuous efforts we were unable to find Cx26 immunoreactivity in the CB at any stage of postnatal development. Thus, the carotid body could not contribute our the *in vivo* WBP phenotype following ablation of Cx26.

Conversely, Cx26+ cells in which none of the Cre-drivers are active will not yield the observed mutant phenotype either. For example cells, such as those of the (ventral diencephalic) hypothalamus, which are neither positive for Cx26 (Mercier and Hatton, 2001) nor derived from dorsal neural tube or neural crest (and thus unlabelled by the well published neural crest Cre-transgenes we utilized), cannot contribute to a phenotype in the mutants either. Similarly, the locus coeruleus, which is located near the lateral floor of the fourth ventricle, is derived from a rhombomeric part that is never positive for the Wnt1-Cre or P0-Cre transgenes (Aroca et al., 2006; Aroca and Puelles, 2005). Thus Wnt1-cre expression would not elicit Cx26 excision in the locus coeruleus (or its precursor tissue). Furthermore we were unable to find Cx26-IR in the locus coeruleus,

Similarly, as leptomeninges around the brainstem are mesodermal in origin – they are (and remain, as predicted) strongly Cx26+ after neural crest specific Cx26 ablation using the Wnt1-Cre and P0-Cre drivers. So if any (forebrain or midbrain neural crest derived) leptomeningeal Cx26 were to influence the chemosensory control of breathing, PGDS-Cre mediated ablation of Cx26 would have yielded an effect detectable by WBP, yet we failed to see this.

Thus we can confidently exclude the carotid body, leptomeninges, hypothalamus and locus coeruleus from contributing towards the *in vivo* WBP effects in the conditional Cx26 mutants used in this study. A further important point is that we can also exclude developmental loss of the NCC Cx26+ cells as a possible explanation of our results. Our simultaneous Cre-reporter transgene shows that vGFP labelled cells are still present in the mutants. Thus Cx26 does not play a role during the development or establishment of these cells *in vivo*. Instead the phenotype must arise from the selective loss of Cx26 in these cells and the consequent loss of CO_2_-evoked ATP release.

### A novel neural crest derived CO_2_ chemosensory cell type, unique to the caudal brainstem

The identity and origins of the Cx26+ cells relevant to CO2 sensing and breathing control were previously unknown, precluding cell specific gene ablation and functional analysis. As we used the same Cre-transgenes for lineage analysis and Cx26 gene ablation (inside the same animal) this can now be resolved. Although NCCs were not expected to be present in the medulla (Etchevers et al., 2002; Etchevers et al., 2001) (let alone signaling) in this brain region, genetic tracking reveals a discrete NCC population, well hidden within the glia limitans, with unique localization, morphology and function. Being strictly confined to the caudal chemosensitive area, their cellular morphology is consistent with Loeschke’s hitherto untested prediction that central CO_2_ sensing occurs within 200 µm from the ventral medullary surface (Trouth et al., 1973). Unexpectedly this cell type is GFAP+, thus able to hide among GFAP+ astrocytes. It is well known that neural crest does not give rise to (GFAP+) astrocytes in the brain as the latter emerge from more ventral CNS areas during development.

The localization of these NCCs corresponds quite closely to Schlaefke’s area (Schlaefke et al., 1970). Cooling or coagulation of this area abolished the CO_2_ sensitivity of breathing in anesthetized cats (Loeschcke et al., 1979; Schlaefke et al., 1979). Unlike the more rostral and caudal chemosensitive areas, focal acidification to pH 7.0 of this intermediate area reduced rather than increased respiratory activity (Schlaefke et al., 1970). This puzzling observation of more than 4 decades ago now finds its molecular mechanistic explanation in the functional characteristics of the Cx26 hemichannel that we recently revealed: Cx26 closes if made sufficiently acidic (Huckstepp et al., 2010a; Huckstepp et al., 2010b). Such closure of Cx26 due to acidosis would remove the tonic release of ATP from the ventral surface of the medulla (Huckstepp et al., 2010b) and hence reduce respiratory drive (Gourine et al., 2005). Interestingly Schlaefke et al (1970) found that alkalinisation to pH 7.8 increased respiratory activity in this area. This again is consistent with our prior observations of the properties of Cx26: its sensitivity to CO_2_ is increased by alkaline pH (Huckstepp et al., 2010b) and CO_2_-carbamylation is sensitive to pH. The pH-sensitivity of CO_2_-binding to Cx26 therefore provides a unifying molecular concept that suggests that the NCC chemosensors may be the key cellular substrate responsible for the function of the Schlaefke’s area.

### Contribution of Cx26 to the regulation of breathing

Our results establish Cx26 as a molecular transducer for direct central CO_2_ chemoreception in mammals *in vivo*. Deletion of Cx26 showed that this molecule is a major mediator of CO_2_-dependent ATP release in both the rostral and caudal chemosensory areas. For the chemosensory control of breathing it appears that the Cx26-mediated ATP release occurring from the NCCs uniquely found in the caudal area is of particular importance: deletion of Cx26 from these few cells reduced the adaptive changes in tidal volume that occur in response to hypercapnia by over 40%. We note that the magnitude this contribution of direct CO_2_-sensing via Cx26 to the overall adaptive response to hypercapnia in mice corresponds well to the contribution proposed for direct CO_2_ sensing in the chemosensory control of respiration in the cat by Shams (1985).

A consensus regards detection of pH as the sole but indirect mechanism to mediate central (medullary) CO_2_ chemosensitivity (Loeschcke, 1982). This consensus has recently received mechanistic support from the observation that global (i.e. not a cell-type specific) genetic ablation of GPR4 greatly reduces chemosensory responses and that combined unconditional deletion of both GPR4 and TASK2 apparently eliminates chemosensory responses in the few surviving knockout animals (Kumar et al., 2015). How do we therefore reconcile our data pointing to a role for Cx26 with these recent data on the importance of pH sensing?

It is well known that responses to hypercapnia involve both an increase in respiratory frequency and an increase in tidal volume. Indeed focal stimulation of the RTN by CO_2_ microdialysis was previously shown to evoke only an increase in tidal volume (Li et al., 1999). However, at variance with these prior findings, deletion of GPR4 in mice affects mainly adaptive changes in respiratory frequency and has only minor effects on the adaptive changes in tidal volume –see Figure S2A,B of Kumar et al. (2015) -especially at the levels of CO_2_ we have used in this study. Our discovery that the effects of Cx26 on the CO_2_ sensitivity of breathing via the caudal chemosensory region are mediated entirely via changes in tidal volume are thus complementary to the role of GPR4 as seen in a global GPR4 knock out (Kumar et al., 2015). The effects of Cx26 deletion are prominent only over an age window of about 3-7 months, encompassing “middle-age”. The age-dependence of the effects of GPR4/TASK2 deletion have not been reported systematically, but it is conceivable that further complementarity will be apparent when this has been studied and once cell-specific gene ablations of GPR4/TASK2 have been performed in an age-specific fashion, in parallel to our study.

Other evidence supports the role of RTN neurons in the chemosensory signaling pathway. Many RTN neurons express *Phox2b*, a transcription factor, which is implicated in congenital central hypoventilation syndrome (CCHS). CCHS is regarded as a defect of central chemosensitivity, and when the human mutations of *Phox2b* are selectively expressed in the RTN, mice lose all chemosensitivity to CO_2_ leading to substantial postnatal lethality (Ramanantsoa et al., 2011). However, in the mutant mice that do survive, there is a recovery of about 40% of the chemosensitivity as they mature to adulthood. This restoration of the responses to CO_2_ occurs entirely as adaptive changes in tidal volume after about 3 months of age. We can now propose a first mechanistic explanation for this phenomenon. We posit that this recovery of central chemosensitivity in Phox2b mutants is consistent with the maturation of a separate pathway involving Cx26-dependent detection of CO_2_ via the NCCs that we describe in this paper.

Cx26 hemichannels may also be important for the control of breathing in humans. The Cx26 mutation A88V prevents CO_2_-dependent gating of the hemichannel (Meigh et al., 2014). This mutation causes human keratitis ichthyosis deafness (KID) syndrome (Koppelhus et al., 2010). We recently found a KID syndrome infant carrying this mutation who exhibited repeated periods of central apnea suggestive of blunted chemosensory control and altered respiratory drive (Meigh et al., 2014). Such a human mutation would be expected to affect all Cx26+ cell types that sense CO_2_, including blood vessel endothelia. In this paper we have specifically ablated Cx26 only in certain neural cell types. Future ablations of Cx26 in vasculature might also yield similar phenotypes to those of the KID syndrome patients..

### Postnatal development of chemosensory modulation of breathing

Surprisingly, mutant phenotypes associated with Cx26 deletion from NCCs in the caudal area only become apparent between 3 and 9 months. Genetic lineage labelling enabled us to associate the time course of this phenotype with the clonal expansion, colonization and later loss of these Cx26+ NCCs. This questions the frequent assumption that all key sensor systems are in place at birth (Huckstepp and Dale, 2011). Our study unveils transient anatomical changes within the adult period that are of physiological importance. We advocate extending the current subdivision of postnatal respiratory CO_2_ chemosensitivity into three phases (juvenile, sexually mature and old adult) with a fourth period of ‘middle-age’, when care for the young is central (Figure 8). It will be interesting if middle-age-related changes in chemosensitivity in humans can be identified and mapped onto hitherto orphan diseases affecting awake patients (Nogues et al., 2002). Interestingly, obesity hypoventilation syndrome (OHS) largely affects patients in middle-age and involves central sleep apnoea in addition to obstructive sleep apnoea (Mokhlesi, 2010) and is a potential candidate for a Cx26-dependent process. Once the exact rhombomeric origins of these NCCs will be identified, focused searches for genes involved become feasible: any gene or epigenetic process negatively affecting the postnatal ontogeny (migration, integration, survival, function) of these few but essential NCCs could be aetiological for tidal volume related breathing phenotypes among the host of apneas and hypopneas known in humans, many of which only emerge in middle age.

### Unexpected neurocristopathies – a link to Chiari 1 malformation

The extent of the reduction of the change in tidal volume in response to hypercapnia in the Wnt1:Cx26-KO mice is remarkably congruent with this prediction. Surprisingly, the primary CO_2_ sensors relevant for breathing are neither neurons nor GFAP+ astrocytes, but instead are superficial GFAP+ NCCs carrying Cx26. Respiratory defects are known in neurocristopathies (Chen and Keens, 2004; Dauger et al., 2003; Gaultier et al., 2005; Poceta et al., 1987), but were hitherto only attributed to peripheral defects of the NCC-derived carotid body in which we could not find Cx26+ cells (see Figure 1, Supplement 2 and Table 3). Thus, the carotid body is unlikely to contribute substantially to our observed WBP phenotypes. Cx26 is essential for CO_2_-dependent ATP release in the isolated medullary slice -a recognized key event of central chemoreception. While NCCs form all sensory neurons of the peripheral nervous system (Le Douarin and Kalcheim, 1999), this is the first extension of the neural crest ‘sensor concept’ to the brain parenchyma that we are aware of.

The Arnold-Chiari I-malformation which has a frequency of 1% in the human population affects the medulla within the posterior cranial fossa (Speer et al., 2003). We previously identified it as a neurocristopathy on the basis of genetic lineage analysis in the cranial base skeleton (Matsuoka et al., 2005). Chiari 1 patients commonly experience breathing defects (apneas, hypopneas) (Dauvilliers et al., 2007; Ferre Maso et al., 2014; Henriques-Filho and Pratesi, 2008; Kijsirichareanchai et al., 2014). Based on our findings it is a plausible hypothesis that the NCCs we have identified as chemosensors may be compromised in these patients. A significant part of the Chiari1 patient population is only diagnosed in the 2^nd^ and 3^rd^ decade (Milhorat et al., 1999), a time in human ontogeny that is comparable to the middle-age phase of child-rearing in which Cx26 is affecting adult mouse breathing. We hypothesize and predict that the ventrally located CO_2_-sensing cells identified in this study might become mechanically damaged by posterior fossa crowding or lost due to primary genetic defects within these cells. The caudal chemosensory region we identified is in close proximity to the basioccipital, the end of the clivus and the craniocervical junction, which is severely affected in Chiari 1 patients. Our hypothesis is more directly testable by studying whether hypercapnic stimuli give rise to blunted changes in tidal volume in a significant fraction of adult Chiari I patients. If our hypothesis is supported, then breathing tests might become part of the early diagnostic toolkit for diagnosing and predicting adult-onset Chiari I, a debilitating condition that becomes symptomatic without prior warning. As ATP actions are accessible to pharmacological manipulation, new interventions can also be considered.

### Living in burrows: a carer’s breath to protect the young

Why should awake adults in middle age exhibit increased breathing responses to 6% CO_2_ compared to younger animals? Rodents live in burrows and nests with small altricial young, a feature inherited from our small synapsid ancestors, animals such as *Thrinaxodon* that are known to have lived in burrows already (Damiani et al., 2003). In the wild, hypothermia is still the top mortal threat to small mammalian pups (Berry and Bronson, 1992). Adaptations such as huddling in small burrows counter this hypothermia but increase local hypercapnia, often significantly exceeding 6% CO_2_ in the inspired air (Boggs et al., 1984; Shams et al., 2005; Williams and Rausch, 1973). For the pup-rearing mother heat loss is less critical to her own survival due to her higher metabolic rate, lower surface area to volume ratio and presence of insulating fur. When adults enter the burrow, CO_2_ levels and ambient temperature rise (Shams et al., 2005; Williams and Rausch, 1973). We speculate that this local hypercapnia triggers a Cx26-dependent increase in the depth of parental breathing. Such increased expiration may turn out to be an ancient behavioural adaptation to enhance heat transfer from carers to their pups. Once all circuit elements are identified and become functionally testable the sophisticated mechanisms linking adult respiratory control, metabolic energy control, eusocial behavior (Smorkatcheva and Lukhtanov, 2014) and their evolutionary and biomedical implications will become visible and therapeutically accessible.

## Materials and Methods

### Ethics

All procedures on animals were evaluated by the Animal Welfare and Ethical Review Board of the University of Warwick and carried out in strict accordance with the Animals (1986) Scientific Procedures Act of the UK under the authority of PPL 80/2555.

### Mouse strains

The floxed Cx26 mice (Cohen-Salmon et al., 2002) were obtained from the European Mouse Mutant Archive. The GFAP-cre mice (Tg(Gfap-cre)73.12Mvs, Stock No: 102886) were obtained from Jackson Labs. The P0-cre mice (C57BL/6J-Tg(P0-Cre) 94 Imeg mouse strain, CARD ID 148) were provided by CARD, IRDA, Kumamoto University, Japan (Yamauchi et al., 1999). The Wnt1-cre mice were obtained from Jackson labs (Stock No. 003829). The PGDS-cre mice were obtained from Dr Marco Giovannini (House Research Institute, Los Angeles) (Kalamarides et al., 2011). The R26R-Confetti mice (Stock No: 013731) (Snippert et al., 2010) were obtained from Jackson Labs.

### Recombinase reporter with membrane GFP targeting

In order to visualize all of the cells once expressing Cre and their descendants, we made a Cre reporter transgenic mouse.

1. VenusGFP is developed from YFP with faster and more efficient maturation (gift from Dr. Atsushi Miyawaki, RIKEN, Japan. Nagai, T). VenusGFP was without ATG was made by PCR and put behind the lox2272 and was made in frame with GAP43 and lox2272.
2. To show the whole cell shape, membrane localization signals, N-term 20 amino acids of mouse GAP43, corresponding to AA 150-209 of NM_008083.2, were added in front of the lox2272 3XpolyA sequences. GAP43 was initially chosen using tissue culture experiments investigating the best, most ubiquitous membrane localization in the largest variety of different cell types.
3. Human UqC promoter is ubiquitous promoter that can drive reliable gene expression across different cell types and tissues. The promoter sequence is from −1to −1540 counting from the starting site of the coding sequence. (AGGCTCAGGGAGGTTGAAGGGGGCTGAGCAAAGGAAGCCCCGTCATTACCTCAAATGTGACCCAAAAATAAAGACCCGTCCATCTCGCAGGGTGGGCCAGGGCGGGTCAGGAGGGAGGGGAGGGAGACCCCGACTCTGCAGAAGGCGCTCGCTGCGTGCCCCACGTCCGCCGAACGCGGGGTTCGCGACCCGAGGGGACCGCGGGGGCTGAGGGGAGGGGCCGCGGAGCCGCGGCTAAGGAACGCGGGCCGCCCACCCGCTCCGGGTGCAGCGGCCTCCGCGCCGGGTTTTGGCGCCTCCCGCGGGCGCCCCCCTCCTCACGGCGAGCGCTGCCACGTCAGACGAAGGGCGCAGCGAGCGTCCTGATCCTTCCGCCCGGACGCTCAGGACAGCGGCCCGCTGCTCATAAGACTCGGCCTTAGAACCCCAGTATCAGCAGAAGGACATTTTAGGACGGGACTTGGGTGACTCTAGGGCACTGGTTTTCTTTCCAGAGAGCGGAACAGGCGAGGAAAAGTAGTCCCTTCTCGGCGATTCTGCGGAGGGATCTCCGTGGGGCGGTGAACGCCGATGATTATATAAGGACGCGCCGGGTGTGGCACAGCTAGTTCCGTCGCAGCCGGGATTTGGGTCGCGGTTCTTGTTTGTGGATCGCTGTGATCGTCACTTGGTGAGTAGCGGGCTGCTGGGCTGGCCGGGGCTTTCGTGGCCGCCGGGCCGCTCGGTGGGACGGAAGCGTGTGGAGAGACCGCCAAGGGCTGTAGTCTGGGTCCGCGAGCAAGGTTGCCCTGAACTGGGGGTTGGGGGGAGCGCAGCAAAATGGCGGCTGTTCCCGAGTCTTGAATGGAAGACGCTTGTGAGGCGGGCTGTGAGGTCGTTGAAACAAGGTGGGGGGCATGGTGGGCGGCAAGAACCCAAGGTCTTGAGGCCTTCGCTAATGCGGGAAAGCTCTTATTCGGGTGAGATGGGCTGGGGCACCATCTGGGGACCCTGACGTGAAGTTTGTCACTGACTGGAGAACTC GGTTTGTCGTCTGTTGCGGGGGCGGCAGTTATGGCGGTGCCGTTGGGCAGTGCACCCGTACCTTTGGGAGCGCGCGCCCTCGTCGTGTCGTGACGTCACCCGTTCTGTTGGCTTATAATGCAGGGTGGGGCCACCTGCCGGTAGGTGTGCGGTAGGCTTTTCTCCGTCGCAGGACGCAGGGTTCGGGCCTAGGGTAGGCTCTCCTGAATCGACAGGCGCCGGACCTCTGGTGAGGGGAGGGATAAGTGAGGCGTCAGTTTCTTTGGTCGGTTTTATGTACCTATCTTCTTAAGTAGCTGAAGCTCCGGTTTTGAACTATGCGCTCGGGGTTGGCGAGTGTGTTTTGTGAAGTTTTTTAGGCACCTTTTGAAATGTAATCATTTGGGTCAATATGTAATTTTCAGTGTTAGACTAGTAAATTGTCCGCTAAATTCTGGCCGTTTTTGGCTTTTTTGTTAGACA)
4. A pair of loxP sites is the substrate of Cre recombinase. Lox2272 is a mutant of loxP site with high recombination efficiency with the identical mutant but never with the wild-type loxp which can avoid recombination between widely used loxp and lox2272 if two floxed sited exist in the same cell on different chromosomes. 3 SV40 polyA sequences were put between two lox2272 sites to stop the translation of VenusGFP prior to Cre-recombination. Lox2272 were assembled in the inverted orientation to avoid the stop codons in the left lox2272 site after recombination.
5. Two copies of chicken β-globin HS4 insulator were added to both ends of the transgene (pink) to insulate the construct from external enhancers and silencing chromatin.

For every step, all of the fragments were sequenced. The whole construct was tested in the HEK 293 cells prior to transgenesis.

### Genotyping of transgenes

The genotyping was done by PCR. All of the Cre lines were genotyped with primers: 5’-GCTGGTTAGCACCGCAGGTGTAGAG-3’ and 5’-CGCCATCTTCCAGCAGGCGCACC-3’. For Cx26 flox/flox, the primers are: 5’-CTTTCCAATGCTGGTGGAGTG-3’ and 5’-ACAGAAATGTGTTGGTGATGG-3’ and products are 290bp for wildtype and about 400bps for floxed copy. The primers for XZGFP are: 5’-GCACGACTTCTTCAAGTCCGCCATGCC-3’ and 5’-GCGGATCTTGAAGTTGGCCTTGATGCC-3’ with 243bps product. The confetti mouse was genotyped according to The Jackson Laboratory protocol for stock 013731.

### Immunohistochemistry

For frozen fixed sections of mice harbouring the mvGFP reporter, we perfused with 4% PFA in PBS. Brains were dissected out and fixed further in 4% PFA for 2 hours to overnight and cryo-protected in 30% sucrose overnight. The brains were embedded in OCT and sectioned at 12-17µm. The slides were blocked with 20% FCS in PBS with 0.2% Triton. The primary antibodies were diluted in PBS as below: chicken anti-GFP (Abcam; 1:400-1:800), mouse anti-Cx26 (Invitrogen; 1:200), rabbit anti-Cx26 (Invitrogen; 1:200), rabbit anti-GFAP (Abcam 1:200), cy3-conjugated mouse anti-GFAP (Abcam 1:400) and goat anti-GFAP (Abcam 1:200) and incubate overnight at 4C. Different Alexa dye- (Invitrogen 1:600) or cy3-cy5-conjugated secondary antibodies were added and incubated at room temperature for 1 hour. Finally the sections were stained with DAPI. Confocal images were obtained on a Leica SP5 or Zeiss710 confocal microscope.

### Whole body plethysmography

CO_2_-induced respiratory responses in conscious mice were recorded using whole body plethysmography as descibed in detail previously (Trapp et al., 2008; Trapp et al., 2011). A plethysmograph was constructed from a plexiglass box (405 ml) equiped with a tight fitting lid, a pressure transducer to detect the respiratory movements of the mouse, and gas inlets and outlets to permit gas flow through the chamber. A humidified mixture of O_2_ (∼20%), N_2_ (80%) and CO_2_ (0-6%) flowed through the chamber at a rate of 1 l.min^-1^. The amount of O_2_ and CO_2_ in the mixture, just prior to entry into the chamber, was measured by a gas analyzer (Hitech Intruments, GIR250 Dual Sensor Gas analyzer). The plethysmograph was calibrated on each use by injection of defined volumes of air. The output of the pressure transducer was amplified, filtered and then recorded to a hard disk via a CED micro1401 interface and Spike2 software.

One experimenter performed the plethysmography, while a second, blind to the genotype of the mice involved, performed the analysis of the records offline at a different time and location. A mouse was placed in the chamber and allowed at least 20 minutes to acclimatize, before being exposed successively to 3% and 6% CO_2_ each for 5 minutes. The plethysmography recordings were analyzed in two ways using Spike2 software. This software was used to trigger a cycle-by-cycle average of the plethysmography wave form for each breathing cycle, giving a measure of tidal volume and respiratory rate (from the mean time between waveform peaks). This method required selection of sequences in the plethysmography trace in which the mouse was judged to be quiescent and exhibiting eupneic breathing as opposed to exploratory sniffing. Tidal volume was normalized to body weight.

An alternative and quicker method of analysis, that did not require data selection, was to take the last 20-60s of data under each condition (the precise amount being determined by the longest period free from gross movement artefact, and was kept consistent for all three conditions). FFT analysis (using the Spike2 software) was then performed to create a power spectrum for the breathing movements. This spectrum contained peaks at different frequencies corresponding to eupneic breathing, as well as other higher frequencies that correspond to exploratory sniffing. The power spectrum gave a rapid measure of both tidal power and respiratory rate. Comparison of the two methods over several independent replications indicated that they gave similar results. Analysis of periodic signals is best performed in the frequency domain –and has long been used to analyze delta rhythms during different sleep wake states for example. Its use to analyze breathing movements is less common but not unprecedented (Wertheim et al., 2009). Tidal power is clearly related to the more commonly measured tidal volume. Tidal volume however measures just the peak-to-peak changes in volume, whereas tidal power measures the integral of the breathing movement and is thus a more robust measure. As the FFT analysis was more rapid to implement, and gave a better estimate of respiratory rate, this method was adopted for all analysis. To further validate our analysis, one batch of recordings was also sent to AVG (blind to phenotype), for independent analysis. Tidal power was normalized to body weight.

The deletion of the Cx26 gene was controlled by the presence or absence of the Cre recombinase (maintained as a hemizygote and expressed under the control of a specific promoter). Thus Cre^+/-^ and Cre^-/-^ littermates were effectively the knockout and littermates for comparison. These knockouts and littermates were experimented in an interleaved design with the breathing of both genotypes being measured on the same day and under as near identical conditions as possible. The data from individual trials were then plotted as cumulative probability distributions with each point representing a measurement from a single mouse. The medians of these distributions were compared via the Mann Whitney U test. One-tailed statistics were used as there was a clear *a priori* prediction that deletion of Cx26 should either reduce CO_2_ sensitivity or have no effect.

### Biosensor measurements

Mice of various genotypes were humanely sacrificed in accordance with the UK 1986 Animals (Scientific Procedures) Act by an overdose of inhalation anaesthetic. Using previously described methods (Huckstepp et al., 2010b), the brain stem was carefully isolated and a horizontal slice prepared taking care to preserve the leptomeninges and avoid damage to the ventral surface of the slice. The experimenter was blind to genotype.

Following isolation, the tissue slice was left for 30 minutes to recover in standard artificial cerebrospinal fluid (aCSF) at 33^°^C under constant superfusion at a rate of approximately 6 ml.min^-1^. The following solutions were used:

#### Standard aCSF

124 mM NaCl, 3 mM KCl, 2 mM CaCl_2_, 26 mM NaHCO_3_, 1.25 mM NaH_2_PO_4_, 1 mM MgSO_4_, 10 mM D-glucose saturated with 95% O_2_/5% CO_2_, pH 7.5, PCO_2_ 35 mmHg.

#### 80mM HCO_3_^-^ aCSF

70 mM NaCl, 3 mM KCl, 2 mM CaCl_2_, 80 mM NaHCO_3_, 1.25 mM NaH_2_PO_4_, 1 mM MgSO_4_, 10 mM D-glucose, saturated with 12% CO_2_ (with the balance being O_2_) to give a pH of 7.5 or and a PCO_2_ of 70 mmHg.

All chemicals and compounds were from Sigma, unless otherwise indicated. Glycerol (2 mM) was added to all solutions to enable operation of the ATP biosensor.

ATP and null biosensors (Huckstepp et al., 2010b; Llaudet et al., 2005) were obtained from Sarissa Biomedical Ltd (Coventry). These biosensors have a permselectivity layer that greatly enhances their selectivity for ATP versus non-specific electroactive interferents. They were used in conjunction with a Duostat ME200+ (Sycopel International Ltd). The biosensors were bent so that the sensing portion (0.5 mm in length) could be laid flat against the surface of the slice. Dual simultaneous recordings were made with the ATP and null biosensors placed at both the rostral and caudal chemosensing areas. The null biosensors lack the ATP-sensing enzymes and act as a control for any non-specific signals (Huckstepp et al., 2010b). For every experiment the sensors were calibrated at the end of each recording, and tested against a potential interferent (5HT) to assess selectivity and tested against aCSF with high PCO_2_ to test whether this treatment affected the null and ATP biosensors equally. All records are presented as the differential signal between the ATP and null biosensors. The ATP release data was analyzed offline and blind to genotype. Only the release resulting from the first episode of hypercapnia was measured. The data from individual slices were then plotted as cumulative probability distributions with each point representing measurement of ATP release from a single slice. Mann Whitney U tests were used to compare the release of ATP from slices derived from the wild type and Cx26 mutant mice.

## Acknowledgements

We thank Sam Dixon and Ian Bagley for making available Cx26 conditional mutants, H. Clevers for providing the Confetti-reporters via the Jackson lab, K. Yamamura for providing P0-Cre mice, M. Giovannini for providing the PGDS-Cre mice and M. Sofroniew for assisting in the choice of the GFAP-Cre line. GK thanks Mark S. Kane (Column of Hope) and ConquerChiari for financial support. This work was funded by the MRC (awarded to ND and GK), a ConquerChiari project grant to GK, a Wellcome Trust programme grant awarded to GK, a Wellcome Trust Senior fellowship to AVG and a University of Warwick Chancellor’s PhD fellowship to JMJL.

## Author contributions

ND and GK conceived and directed this study. ND supervised and performed the physiology including data analysis, GK the lineage analysis/genetic aspects. XZ performed genetic experiments, lineage analysis, histology, imaging and plethysmography, JZ performed biosensor measurements, AVG supported various physiological aspects and data analysis, JMJL performed image analysis. ND and GK wrote the paper with input of the other authors. All authors commented on the manuscript.

---

**Figure 1 Supplement 1.**
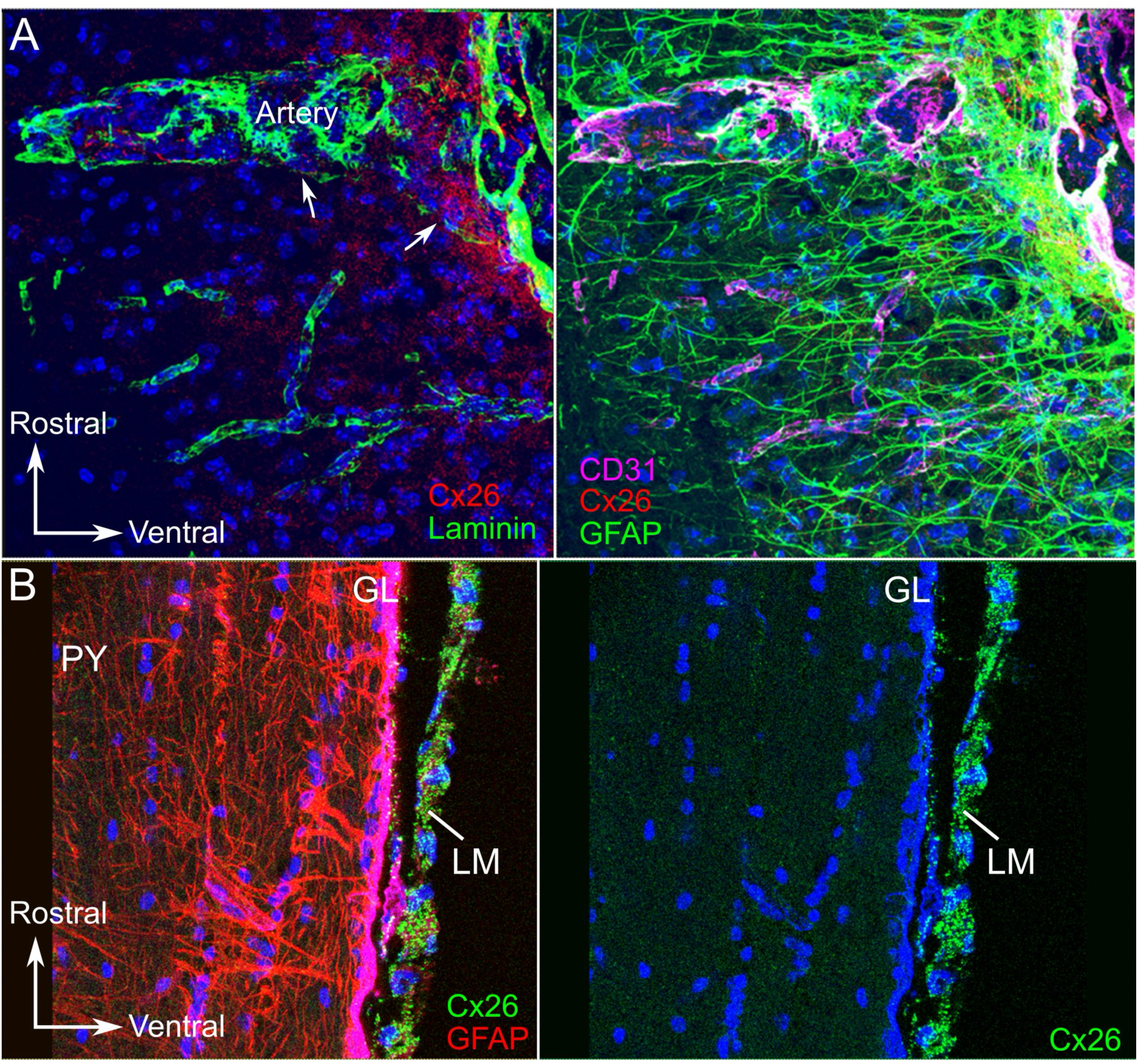
Molecular architecture of the rostral chemosensing region of wild type mouse 4 months old in the vicinity of large vessels (anterior inferior cerebellar artery, AICA). A) Cx26 localization (red) on CD31+ endothelial cells (magenta) of penetrating artery. The basal lamina (laminin, green) is fenestrated at arterial entry points and Cx26-IR is specifically distributed within these fenestrations which would facilitate signaling interactions between Cx26+ arterial vasculature and brain parenchyma (white arrows). B) Outside of the chemosensing regions, Cx26-IR is not observed in the parenchyma. Brain parenchyma and glia limitans of non-chemosensing regions such as the pyramidal tracts (PY) adjacent to the chemosensing regions are not Cx26-immunoreactive. Note the ubiquitous CX26-IR in leptomeninges (LM).

**Figure 1 Supplement 2.**
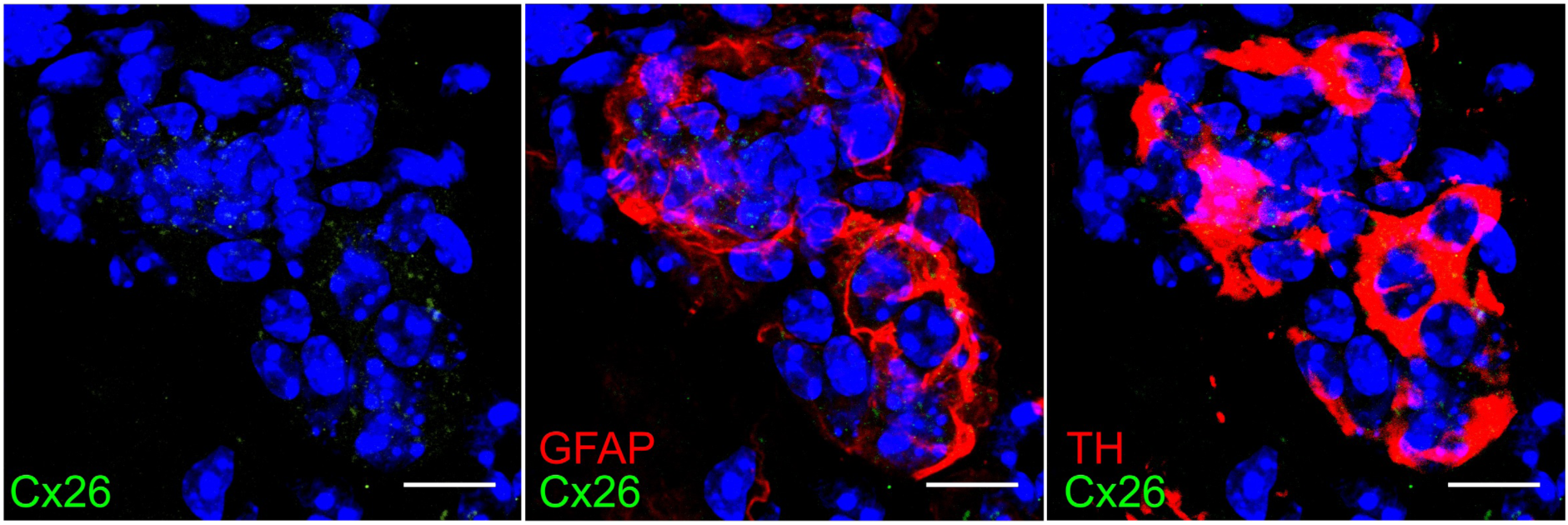
Lack of Cx26 in carotid body. Single optical sections of carotid body stained for Cx26, GFAP and tyrosine hydroxylase (TH). Scale bar 10 µm.

**Figure 1 Supplement 3.**
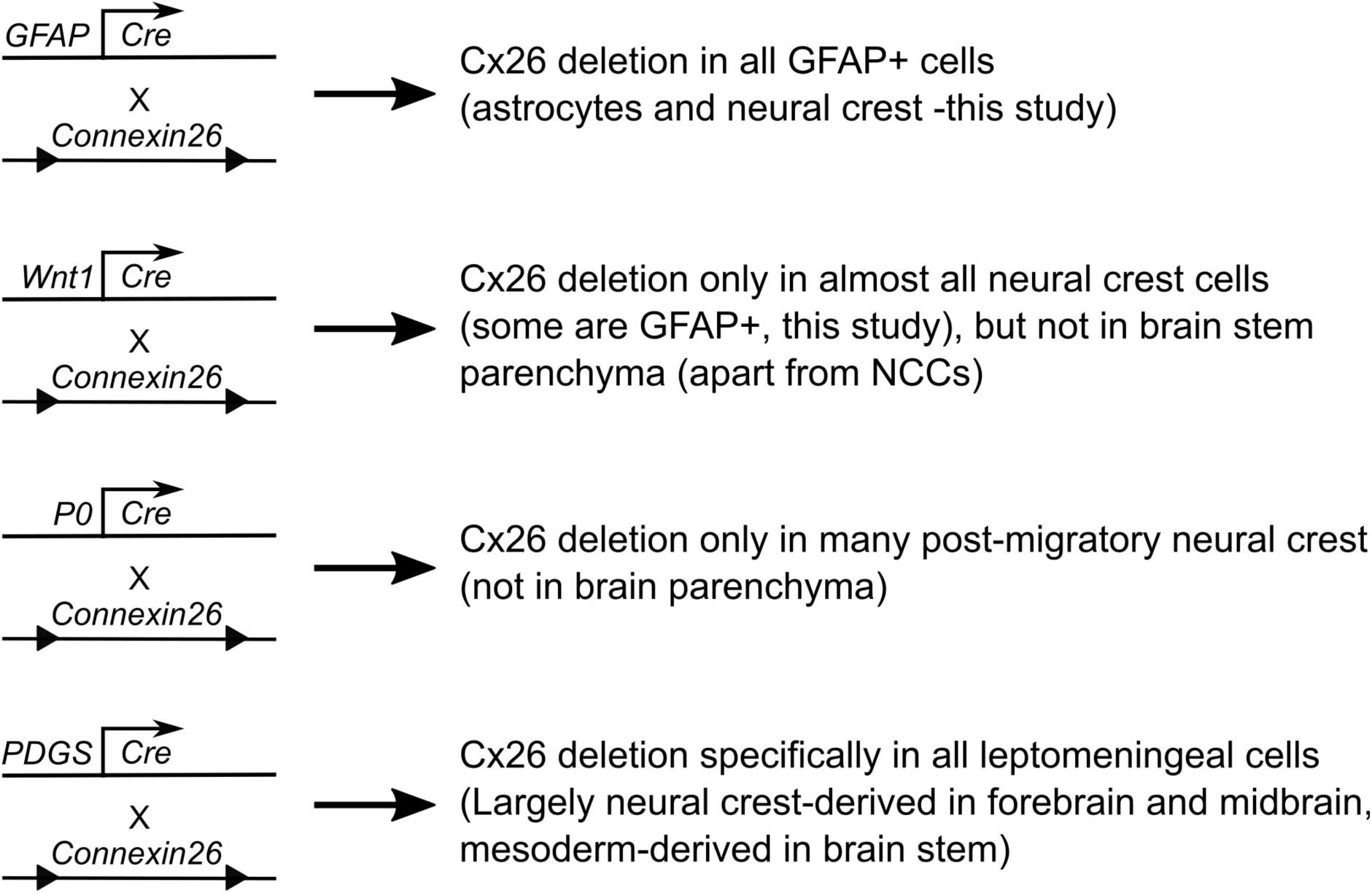
Schematic diagram summarizing genetic strategy for selective deletion of Cx26 from relevant cell populations for this study. Each cross is shown together with target population of cells.

**Figure 5 Supplement 1.**
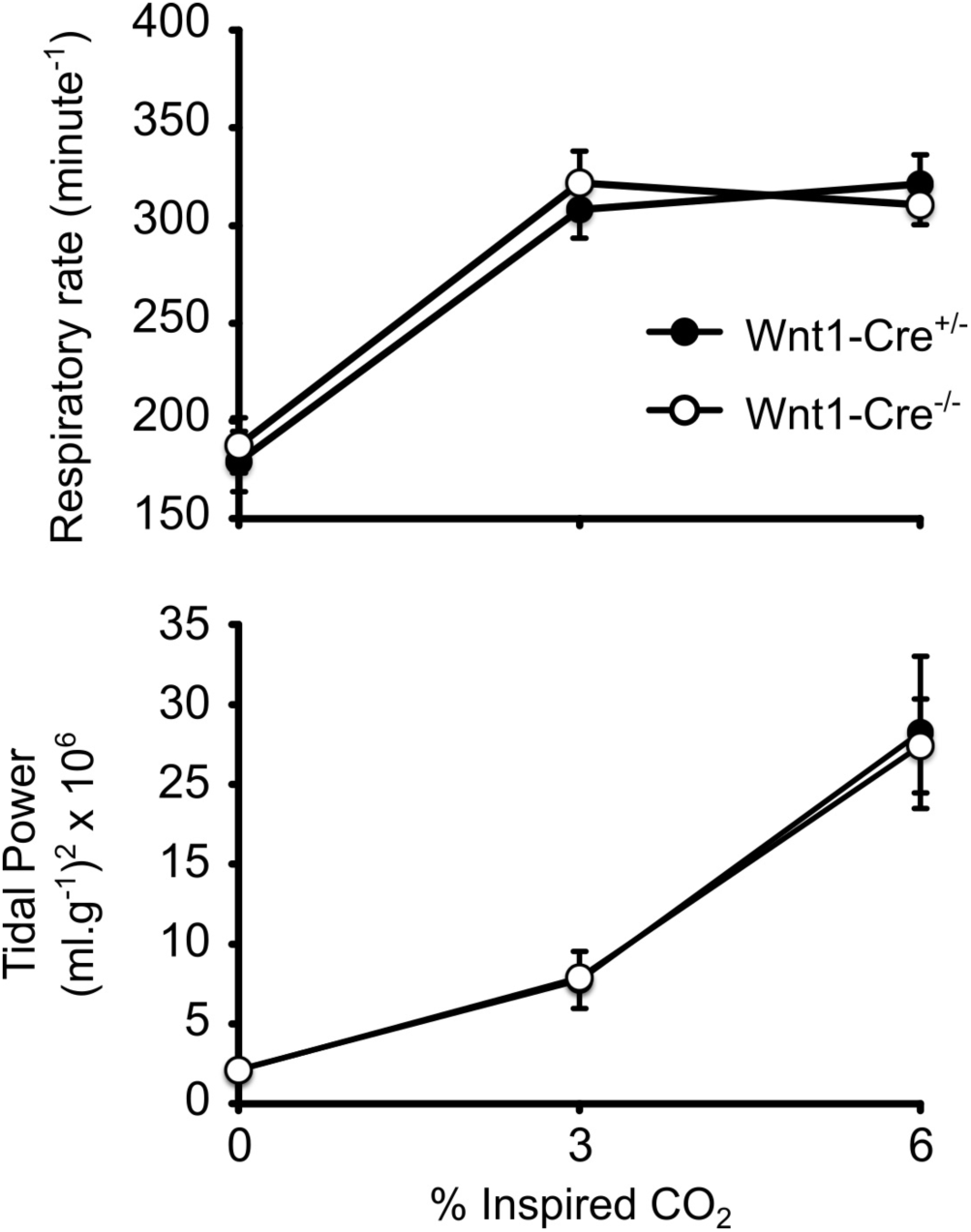
The Wnt1-Cre transgene on a wild type Cx26 background does not affect ventilatory responses to CO_2_. Whole body plethysmography recordings of respiratory rate and tidal power from 10 Wnt1-Cre^+/-^ mice and 10 Wnt1-Cre^-/-^ littermates.

**Figure 5 Supplement 2.**
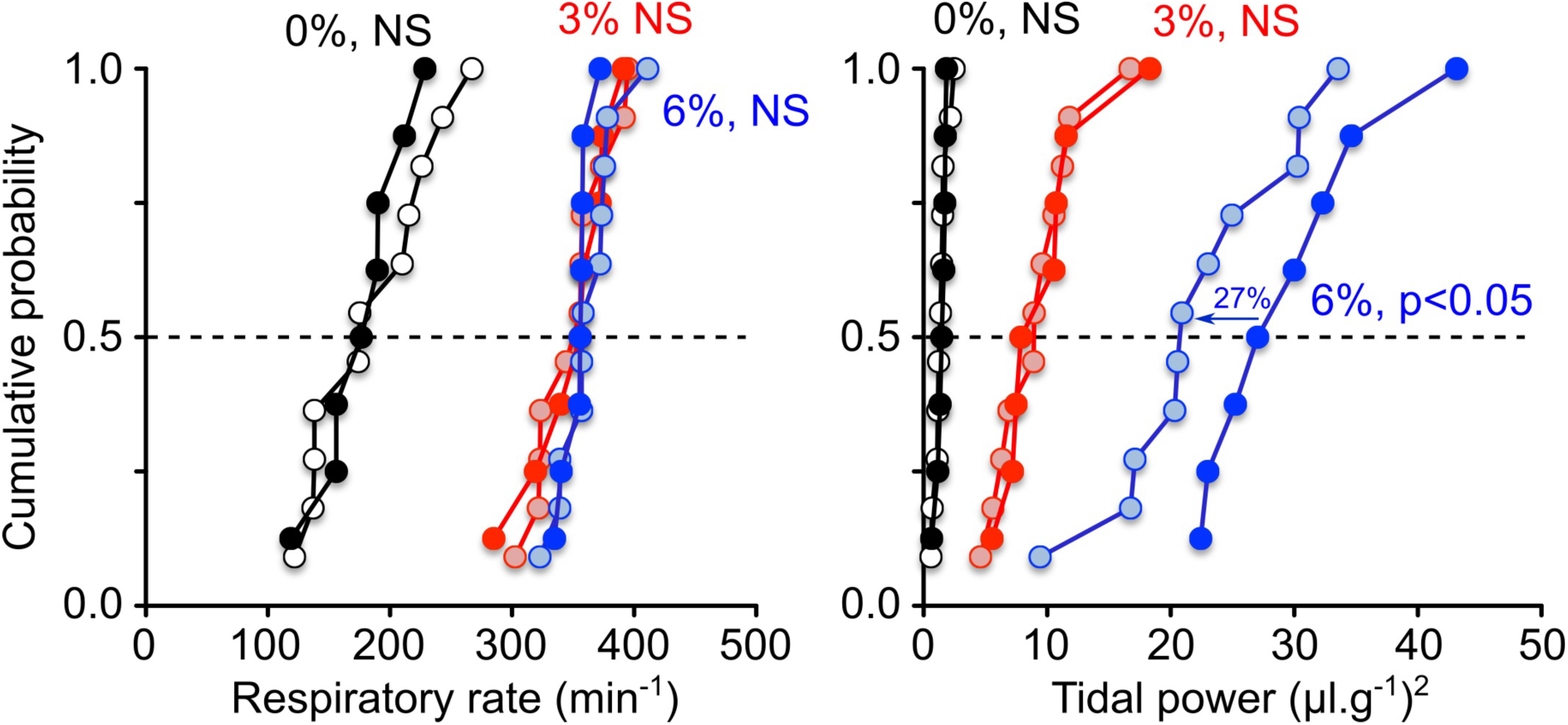
Use of a second genetic marker for neural crest, P0, to delete Cx26 from caudal area causes breathing phenotype. Cumulative probability plots of respiratory rate and tidal power for P0:Cx26-WT and P0:Cx26-KO mice at 0 (black circles), 3 (red circles) and 6% (blue circles) inspired CO_2_ –the lighter filled circles represent the knockout mice at each level of CO_2_. There is no difference in respiratory rate between wild type and knockout mice at any level of inspired CO_2_. The tidal power of breathing for the knockout mice is significantly less than the wildtype mice at 6% inspired CO_2_, but no different at any other level of inspired CO_2_.

**Figure 5 Supplement 3.**
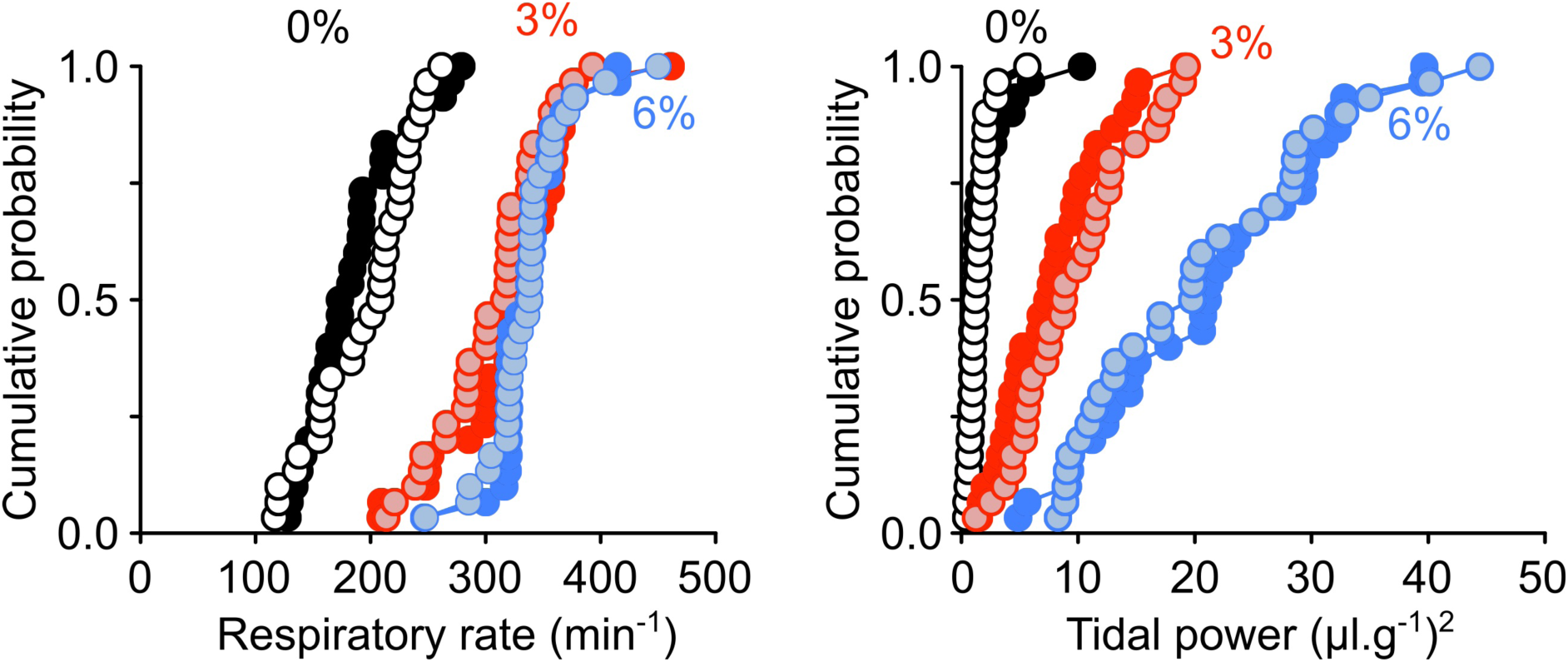
Deletion of Cx26 specifically from the leptomeninges does not alter ventilatory responses to CO_2_. Cumulative probability plots of respiratory rate and tidal power from 30 PGDS:Cx26-WT and 30 PGDS:Cx26-KO mice at 0 (black circles), 3 (red circles) and 6% (blue circles) inspired CO_2_ –the lighter filled circles represent the knockout mice at each level of CO_2_.

